# Oral Hsp90 inhibitor, SNX-5422, attenuates SARS-CoV-2 replication and dampens inflammation in airway cells

**DOI:** 10.1101/2021.02.23.432479

**Authors:** Ria Goswami, Veronica S. Russell, Joshua J. Tu, Philip Hughes, Francine Kelly, Stephanie N. Langel, Justin Steppe, Scott M. Palmer, Timothy Haystead, Maria Blasi, Sallie R. Permar

**Affiliations:** Duke Human Vaccine Institute, Duke University Medical Center, Durham, North Carolina, USA; Department of Pharmacology and Cancer Biology, Duke University Medical Center, Durham, North Carolina, USA; Division of Pulmonary, Allergy and Critical Care Medicine, Department of Medicine, Duke University Medical Center, Durham, North Carolina, USA; Division of Infectious Diseases, Department of Medicine, Duke University Medical Center, Durham, North Carolina, USA; Department of Pediatrics, Duke University Medical Center, Durham, North Carolina, USA

**Keywords:** COVID-19, SARS-CoV-2, Hsp90, Post-exposure prophylaxis, anti-inflammatory, host-directed therapy, orally bioavailable

## Abstract

Currently available SARS-CoV-2 therapeutics are targeted towards moderately to severely ill patients and require intravenous infusions, with limited options for exposed or infected patients with no or mild symptoms. While vaccines have demonstrated protective efficacy, vaccine hesitancy and logistical distribution challenges will delay their ability to end the pandemic. Hence, there is a need for rapidly translatable, easy-to-administer-therapeutics, that can prevent SARS-CoV-2 disease progression, when administered in the early stages of infection. We demonstrate that an orally bioavailable Hsp90 inhibitor, SNX-5422, currently in clinical trials as an anti-cancer therapeutic, inhibits SARS-CoV-2 replication *in vitro* at a high selectivity index. SNX-5422 treatment of human primary airway epithelial cells dampened expression of inflammatory pathways associated with poor SARS-CoV-2 disease outcomes. Additionally, SNX-5422 interrupted expression of host factors that are crucial for SARS-CoV-2 replication machinery. Development of SNX-5422 as SARS-CoV-2-early-therapy will dampen disease severity, resulting in better clinical outcomes and reduced hospitalizations.

## INTRODUCTION

Since its emergence in 2019, severe acute respiratory syndrome coronavirus 2 (SARS-CoV-2) rapidly spread worldwide and the disease associated with SARS-CoV-2 infection, Coronavirus disease 2019 (COVID-19), was declared a pandemic by the WHO in March 2020 (Cucinotta and Vanelli, 2020). As of February 9^th^, 2021, there were more than 106 million confirmed infections and >2.3 million deaths, worldwide (WHO Coronavirus Disease (COVID-19) Dashboard, 2020), making this ongoing disease the deadliest pandemic of the 21^st^ century. Clinical manifestations of SARS-CoV-2 infection can range from asymptomatic (Nishiura et al., 2020) and mild upper airway symptoms to severe lower respiratory complications such as pneumonia, acute respiratory distress syndrome (ARDS), and multi-organ system dysfunction (Huang et al., 2020a), leading to respiratory compromise (Giamarellos-Bourboulis et al., 2020) and death (Mehraeen et al., 2020).

There is an ongoing effort to characterize SARS-CoV-2 pathogenesis and identify prophylactic and therapeutic approaches to suppress COVID-19 severity and improve clinical outcomes. Currently used treatment modalities for SARS-CoV-2 infection include repurposed small molecules such as remdesivir (Beigel et al., 2020), immunomodulatory agents such as corticosteroids (Prescott and Rice, 2020), and virus-specific monoclonal antibodies (Zhou et al., 2020b). However, most of these treatment strategies require intravenous infusions, and are administered in moderately to severely ill patients. Moreover, while prophylactic vaccines for SARS-CoV-2 infection have demonstrated robust protection, vaccine hesitancy and logistical distribution challenges, especially in low-to-middle income countries (LMIC), will restrict their ability to quickly end the global pandemic. Thus, there remains an urgent need for rapidly translatable post-exposure prophylactic interventions that can be administered during the early course of infection to prevent severe disease outcomes, hospitalizations, and reduce viral transmissions. Since such interventions will be administered to patients with no-to-mild symptoms, these therapies should be easy to store and administer, and have a wide margin of safety.

Cellular chaperone protein heat shock protein 90 (Hsp90), in addition to maintaining cellular homeostasis (Zhao and Houry, 2005, Taipale et al., 2010), also facilitates proper folding and functionality of virally encoded proteins (Geller et al., 2007). Hence, Hsp90 serves as host determinant for a wide diversity of viruses (Geller et al., 2012, Rathore et al., 2014, Smith and Haystead, 2017), including Coronaviruses (CoVs) such as severe acute respiratory syndrome coronavirus (SARS-CoV) (Li et al., 2004, Li et al., 2020), Middle east respiratory syndrome coronavirus (MERS-CoV) (Li et al., 2020) and SARS-CoV-2 (Emanuel et al., 2020, Li et al., 2020). Recent *in silico* studies have proposed Hsp90 as a potential anti-SARS-CoV-2 target (Iyad et al., 2020). Furthermore, in a murine model of sepsis, Hsp90 inhibition has been associated with dampened systemic and pulmonary inflammation and acute lung injury (ALI) (Chatterjee et al., 2007), the key phenomena associated with poor SARS-CoV-2 clinical outcomes (Bussani et al., 2020, Pandolfi et al., 2020, Blanco-Melo et al., 2020). Hence, a trial was launched to test the efficacy of intravenously administered Hsp90 inhibitor, ganetespib (ADX-1612) against SARS-CoV-2 infection (Aldeyra Therapeutics Press Release, 2020).

SNX-5422 is an orally bioavailable Hsp90 inhibitor (Fadden et al., 2010, Huang et al., 2009), which is currently in clinical trials for solid state malignancies and lymphomas. The toxicity profile of this small molecule has been extensively characterized in humans, and found to be very well tolerated with no reported adverse effects or evidence of immunosuppression (Infante et al., 2014, Rajan et al., 2011). Since Hsp90 can be exploited by viruses for their replication, here, we have evaluated the efficacy of SNX-5422 as a post-exposure prophylactic treatment for SARS-CoV-2 infection and have mapped the interaction of the drug with primary human airway cell transcriptional profile. Our data reveal that SNX-5422, added shortly after infection, suppresses *in vitro* SARS-CoV-2 replication with a high selectivity index, and influences expression of cellular genes associated with SARS-CoV-2 replication and poor disease outcomes, including airway inflammatory pathways. Our findings support further development of SNX-5422 as an oral post-exposure therapeutic for SARS-CoV-2 infection that will reduce the length of viral infectivity and dampen virus-associated inflammatory responses, initiated early in the course of infection. A post-exposure therapy geared towards individuals in the initial phase of infection, will reduce COVID-19 disease severity and hospitalizations and will prevent additional global deaths, while the distribution of prophylactic vaccines ramps up.

## RESULTS AND DISCUSSION

### SNX-5422, an orally bioavailable Hsp90 inhibitor, is a potent suppressor of SARS-CoV-2 replication

Recent *in silico* drug repositioning studies and single-cell transcriptomic analysis have identified Hsp90 as a host dependency factor for SARS-CoV-2 replication (Iyad et al., 2020, Emanuel et al., 2020). Inhibition of Hsp90 was associated with suppression of SARS-CoV-2 replication *in vitro* (Emanuel et al., 2020, Li et al., 2020, Kamel et al., 2020). Here, we sought to determine if an orally bioavailable, highly selective inhibitor of Hsp90 (SNX-5422) (Okawa et al., 2009), currently in clinical trials for cancer therapy (Infante et al., 2014, Rajan et al., 2011), could serve as a post-exposure prophylactic or early therapeutic for SARS-CoV-2 infection. To mimic post-exposure prophylaxis *in vitro*, human lung epithelial cells (Calu-3), and African green monkey kidney cells (Vero E6) (Ogando et al., 2020) were incubated with SARS-CoV-2 strain USA-WA1/2020 for 1hr, and the virus-exposed cells were treated with increasing doses of SNX-5422 or DMSO (drug-vehicle). Using immunostaining and fluorescent imaging of virus-infected Vero E6 cells, we first assessed the antiviral activity of SNX-5422 by comparing proportion of viral nucleocapsid (NP) protein positive cells in drug-treated vs. vehicle-treated cells. Our data indicate that SNX-5422 attenuated intracellular NP expression (Figure 1A) and proportion of NP+ cells (Figure 1B) in a dose dependent manner, with a 50% inhibitory concentration (IC50) of 2.3μM (Table 1). We further evaluated the antiviral activity of a dose range of SNX-5422 by comparing viral RNA shedding in cell supernatants of drug-treated vs. vehicle-treated Vero cells and human lung epithelial cells. SNX-5422 treatment resulted in a reduction of cell-free viral genomic copies (Figure 1C and 1D) with an IC50 of 0.2μM in both cell types (Table 1). Additionally, SNX-5422-treatment reduced cell-free infectious viral titers (Figure 1E and 1F) with an IC50 of 0.38μM in Vero E6 cells and 0.4μM in Calu-3 cells (Table 1).

**Figure 1.**
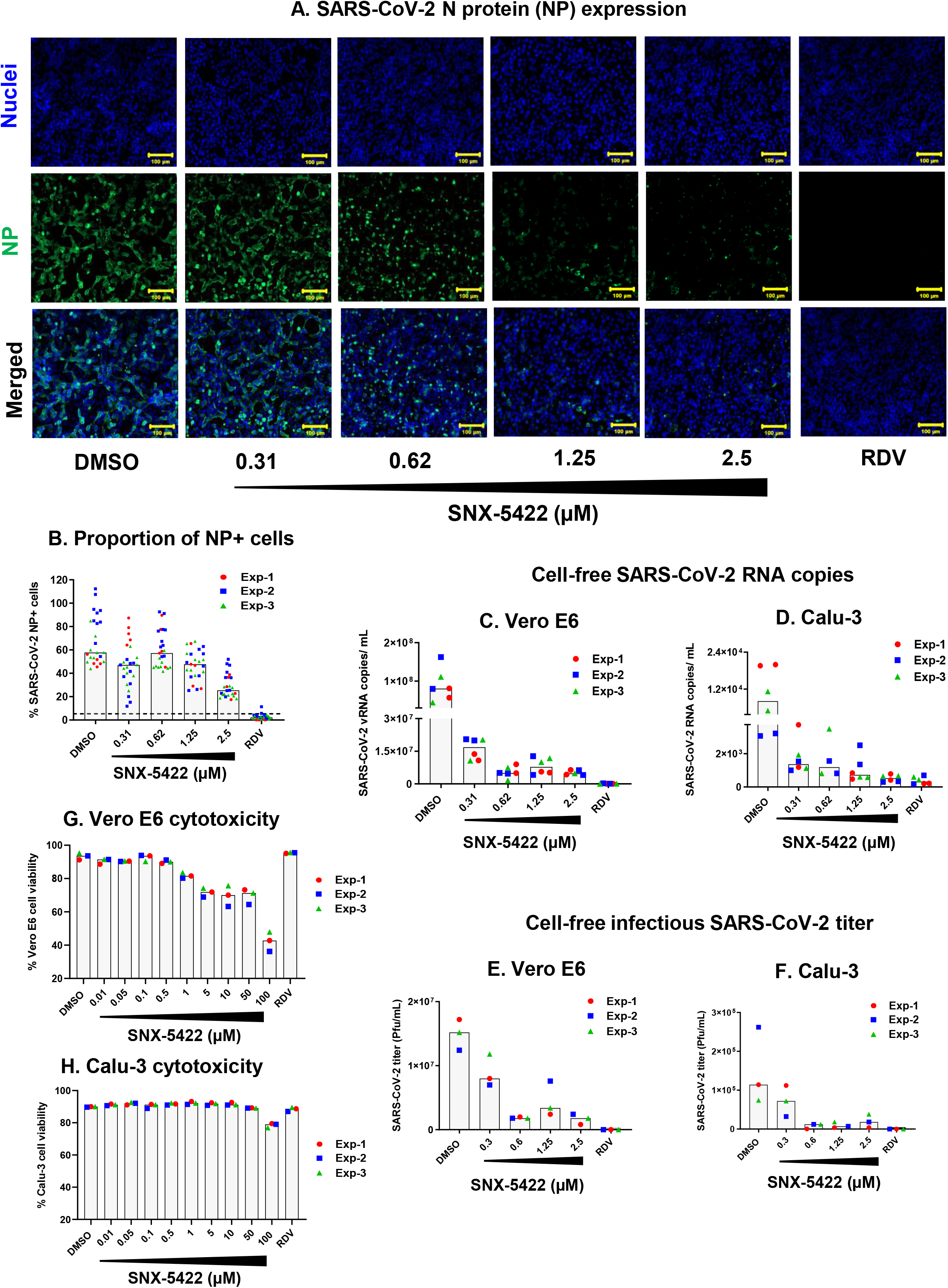
SNX-5422 potently suppresses SARS-CoV-2 replication at non-toxic concentrations. Vero E6 and Calu-3 cells were infected with SARS-CoV-2 strain USA-WA1/2020 and treated with either 0.1% DMSO (drug-vehicle) or 0.31-2.5μM SNX-5422, or 5μM remdesivir (RDV). Both SNX-5422 and RDV were reconstituted in DMSO. After 48 hr post-infection (pi), **(A)** Vero cells were immunostained for SARS-CoV-2 nucleocapsid protein (NP) (green). Nuclei was counterstained (blue) prior to imaging. **(B)** Immunofluorescent images were quantified to evaluate proportions of SARS-CoV-2-NP+ cells. Each symbol and color represent an independent experiment and each data point represents an independent quantified image field. Medians of data points are reported as grey bars. Dotted line represents detection cut off, which is the mean+3 standard deviation (SD) of the percentage of NP+ cells in uninfected controls. Cell-free viral RNA in the supernatant of infected and treated **(C)** Vero-E6 and **(D)** Calu-3 cells were measured by qRT-PCR of the viral N gene. The infectious viral titers of the supernatant of **(E)** Vero E6 and **(F)** Calu-3 were determined by plaque assay. **(G)** Vero E6 and **(H)** Calu-3 cells were treated with either 0.1% DMSO (drug-vehicle) or 0.01-100μM SNX-5422, or 5μM RDV. After 48 hours, proportion of viable cells was evaluated by flow cytometry. Each symbol and color represent an independent experiment and medians of the data points are reported as grey bars.

**Table 1.** Cell-specific SNX-5422 potency and selectivity

| Cytotoxic Concentration (CC) |  |  |  |  |
| --- | --- | --- | --- | --- |
|  | Cell line | Assay | CC50<sup>a</sup><br>(μM) |  |
|  | Vero E6 | Cellular viability | 91.97 |  |
|  | Calu-3 | Cellular viability | >100 |  |
| Inhibitory Concentration (IC) |  |  |  |  |
|  | Cell line | Assay | IC50<sup>b</sup><br>(μM) | SI<sup>c</sup> |
|  | Vero E6 | Cell-free SARS-CoV-2 RNA | 0.2 | 459.85 |
|  |  | Cell-free infectious SARS-CoV-2 titer | 0.38 | 242.03 |
|  |  | Intracellular SARS-CoV-2 N protein expression | 2.3 | 39.99 |
|  | Calu-3 | Cell-free SARS-CoV-2 RNA | 0.2 | >500 |
|  |  | Cell-free infectious SARS-CoV-2 titer | 0.4 | >250 |

<sup>a</sup>CC50: 50% cytotoxic concentration
<sup>b</sup>IC50: 50% inhibitory concentration
<sup>c</sup>SI: Selectivity index=CC50/IC50

Since SNX-5422 inhibits the functionality of Hsp90, (Fadden et al., 2010, Huang et al., 2009), a host chaperone necessary for maintaining cellular homeostasis (Zhao and Houry, 2005, Taipale et al., 2010), we determined the selectivity index of the drug in inhibiting SARS-CoV-2 replication. Uninfected Vero E6 and Calu-3 cells were treated with increasing concentrations of SNX-5422 and the frequency of live cells was monitored by flow cytometry (Figure 1G and 1H). The 50% cellular cytotoxicity (CC50) of SNX-5422 was computed to be 91.97μM and >100μM in Vero E6 and Calu-3 cells, respectively, highlighting the minimal cytotoxic effects of the drug on these two cell types. Consequently, the selectivity index (SI=CC50/IC50) of SNX-5422 in suppressing viral replication was evaluated to be high in both cell types (Table 1). Collectively, these data suggest that treatment with an oral inhibitor of Hsp90, SNX5422, upon SARS-CoV-2 exposure, can potently attenuate viral replication *in vitro*, at a high selectivity index.

### SNX-5422 treatment of primary human tracheobronchial epithelial (TBE) cells alters expression of cellular inflammatory and metabolic pathways

In addition to developing antivirals targeting specific viral proteins, there has been a growing effort in developing therapeutics targeting host proteins exploited by viruses for their replication or implicated in viral pathogenesis (Nitulescu et al., 2020). As such therapeutics can impair normal functioning of host proteins, a clear understanding of their impact on cellular mechanisms is critically important. We therefore characterized the interactions of SNX-5422 with the cellular machinery using a physiologically relevant *ex vivo* human primary tracheobronchial epithelial (TBE) cell model (Mason, 2020). Fully differentiated human TBE cells from three independent donors (Table S1) without known respiratory disease or smoking history were cultured *ex vivo* at the air-liquid interface. The basolateral side of the cell culture was treated with either 0.1% DMSO (drug-vehicle) or 1μM SNX-5422 for 48hrs, followed by bulk RNA sequencing of the total cellular RNA (Figure 2A). Principal component analysis of the read counts revealed that treatment with SNX-5422 formed distinct clusters compared to the control group (Figure S1), highlighting differential expression of cellular genes upon drug treatment. Differentially expressed genes (DEGs) between the control group and the SNX-5422-treated group were defined as protein coding genes with a Log_2_ fold change (FC) ratio > 2 and adjusted p value ≤ 0.001. A volcano plot of the protein coding genes indicated that 1361 genes were differentially expressed between the two groups, with 470 cellular genes being upregulated and 891 cellular genes being downregulated upon SNX-5422 treatment (Figure 2B). To identify cellular genes whose expression was highly regulated upon SNX-5422 treatment of TBE cells, we collated DEGs with a Log_2_ FC >6, for which the average normalized read count across donors and experimental conditions was ≥300 (Table S2). Using this stringent threshold, we identified 16 cellular genes that were downregulated and 4 cellular genes that were upregulated upon drug treatment (Figure 2B). Further categorization of those genes based on functions revealed that SNX-5422-treatment of TBE cells downregulated cytokines and chemokines that serve as chemoattractant for lymphocytes and neutrophils (*CCL20, CXCL1*), regulators of cellular inflammation (*CSF3, S100A8*) and proliferation *(CDC20)*, antigen presentation pathway-associated genes (*HLA-DOA, HLA-DMB*) and genes functioning as solute transporters (*SLC13A2, SLC26A4, SLC5A1, SLC5A5*) (Figure 2B).

**Figure 2.**
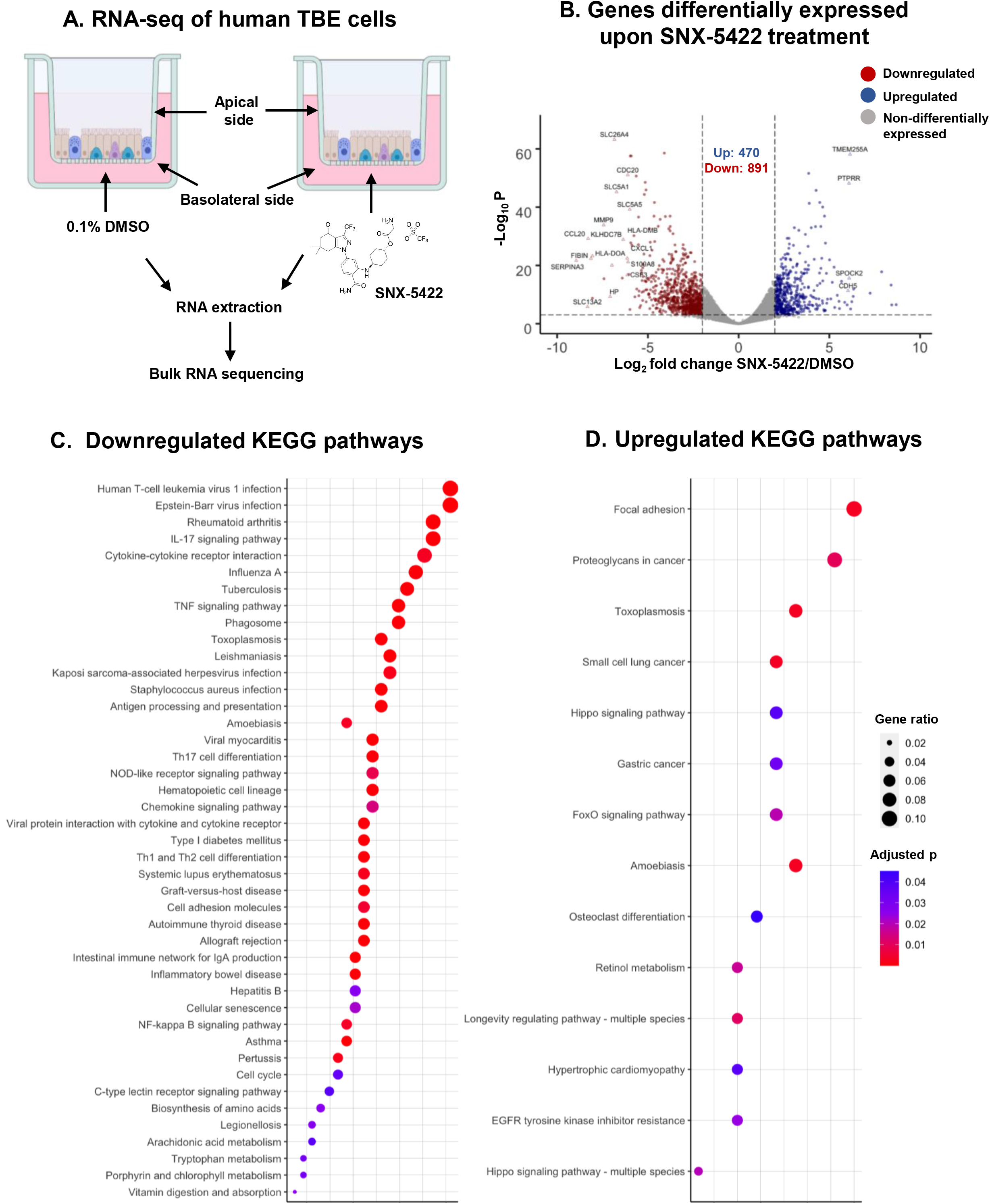
SNX-5422 treatment alters gene expression of human tracheobronchial epithelial (TBE) cells. **(A)** Schematic diagram illustrating treatment of human TBE cells. TBE cells from 3 independent healthy donors were cultured in air-liquid interface and treated with either 0.1% DMSO (drug-vehicle) or 1μM SNX-5422 reconstituted in DMSO, for 48hrs. RNA was then extracted from the cells and bulk RNA sequencing was performed. **(B)** Volcano plot demonstrating protein coding genes altered upon SNX-5422 treatment of human TBE cells. Red, blue and grey indicates downregulated, upregulated and non-differentially expressed genes, respectively. Differentially expressed genes (DEGs) were defined as protein coding genes with a Log_2_ fold change >2 between the drug-treated and control groups and adjusted p value <0.001. Open symbols represent genes with Log_2_ fold change >6 between the drug-treated and control groups. Cellular pathway overrepresentation analysis was performed with DEGs with a normalized average read count across donors and experimental conditions ≥300. **(C)** Downregulated and **(D)** Upregulated Kyoto Encyclopedia of Genes and Genomes (KEGG) pathways with an adjusted p value< 0.05 were reported. Dot size represented gene ratio and color schema represents adjusted p values.

Next, we mapped the cellular signaling pathways that were significantly altered upon SNX-5422 treatment of human TBE cells, by performing a Kyoto Encyclopedia of Genes and Genomes (KEGG) pathway overrepresentation analysis on the DEGs (Figure 2C and 2D). Notably, cellular inflammatory pathways such as IL-17 signaling pathway, cytokine-cytokine receptor interaction, TNF-signaling pathway, Th17 cell differentiation, NOD-like receptor signaling and chemokine signaling pathways were among the top 20 identified pathways that were downregulated by SNX-5422 treatment. In addition, signaling pathways associated with regulation of cell cycle and cellular senescence, and metabolic pathways such as biosynthesis of amino acids and tryptophan metabolism, were also downregulated by SNX-5422 (Figure 2C). Furthermore, treatment with this oral Hsp90 inhibitor resulted in upregulation of key cellular pathways involved in physiological functions such as cellular - adhesion, regulation of apoptosis and cellular proliferation (Figure 2D). Taken together, our data suggest that SNX-5422 alters TBE cell inflammation, cellular proliferation, and metabolism.

### SNX-5422 dampens cellular inflammatory pathways associated with SARS-CoV-2 replication and poor disease prognosis

SARS-CoV-2 infection has been associated with hyper-induction of pro-inflammatory cytokines resulting in multiple organ failure, ALI and ARDS (Song et al., 2020). Interestingly, Hsp90 inhibition dampened systemic and pulmonary inflammation and lung injury in a murine model of sepsis (Chatterjee et al., 2007). Since SNX-5422 exhibited downregulated inflammatory transcriptomic pathways in human TBE cells, we interrogated whether these pathways have been previously associated with SARS-CoV-2 disease progression. To this end, we first selected a gene set whose expression was stringently regulated upon SNX-5422 treatment of human TBE cells (Log_2_ FC >3, total 287 genes) (Table S2). We then performed an extensive literature search in PubMed and non-peer reviewed pre-publication repositories to compare our identified gene set with the cellular genes and pathways previously associated with SARS-CoV-2 replication. To maintain physiological relevance, we restricted our literature search to genes and pathways reported in *ex vivo* SARS-CoV-2-infected primary airway epithelial cells or those identified in blood, bronchioalveolar lavage fluid (BALF) and lung biopsies of SARS-CoV-2 positive patients. Our search identified 55 genes regulated by SNX-5422, which have been associated with SARS-CoV-2 pathogenesis and disease progression. A protein-protein interaction (PPI) network and functional enrichment analysis mapped several of these 55 genes to cellular inflammatory pathways including IL-17 signaling pathway, TNF signaling pathway, cytokine-cytokine receptor interaction and chemokine signaling pathway (Figure 3A). SNX-5422 dampened expression of cellular genes, previously shown to be associated with SARS-CoV-2 -mediated proinflammatory response and hyper-cytokinemia (Figure 3B). Specifically, these inflammatory genes could be broadly categorized into chemokines (*CXCL1* (Coperchini et al., 2020)*, CXCL5* (Coperchini et al., 2020)*, CXCL6* (Blanco-Melo et al., 2020)*, CXCL3* (Blanco-Melo et al., 2020) and *CXCL2* (Zhou et al., 2020a)), chemokine-receptor ligands (*CCL20* (Zhou et al., 2020a)*, CX3CL1* (Nienhold et al., 2020)), cytokines (*IL36G* (Xiong et al., 2020)), interleukins (*IL-19* (Vastrad et al., 2020)*, CXCL8* (Zhou et al., 2020a)*, IL-32* (Blanco-Melo et al., 2020)), tumor necrosis factor alpha-induced protein (*TNFAIP2* (Fagone et al., 2020)), regulators of inflammatory responses (*CSF3* (Nunnari et al., 2020)*, S100A8* (Zhou et al., 2020a)*, S100A9* (Zhou et al., 2020a)*, SCGB3A1* (Mick et al., 2020)*, S100A2* (Xu et al., 2020)) and interferon-induced proteins (*RSAD2 (Zhou et al., 2020a), IFI44L (Mick et al., 2020), IFITM1* (Zhou et al., 2020a)*, MX2 (Lieberman et al., 2020), IFI27 (*Mick et al., 2020)*, ISG15 (Zhou et al., 2020a)*). Collectively, our analysis suggests that early treatment with SNX-5422 may mitigate SARS-CoV-2-mediated hyperinflammation, thereby improving clinical outcomes of COVID-19.

**Figure 3.**
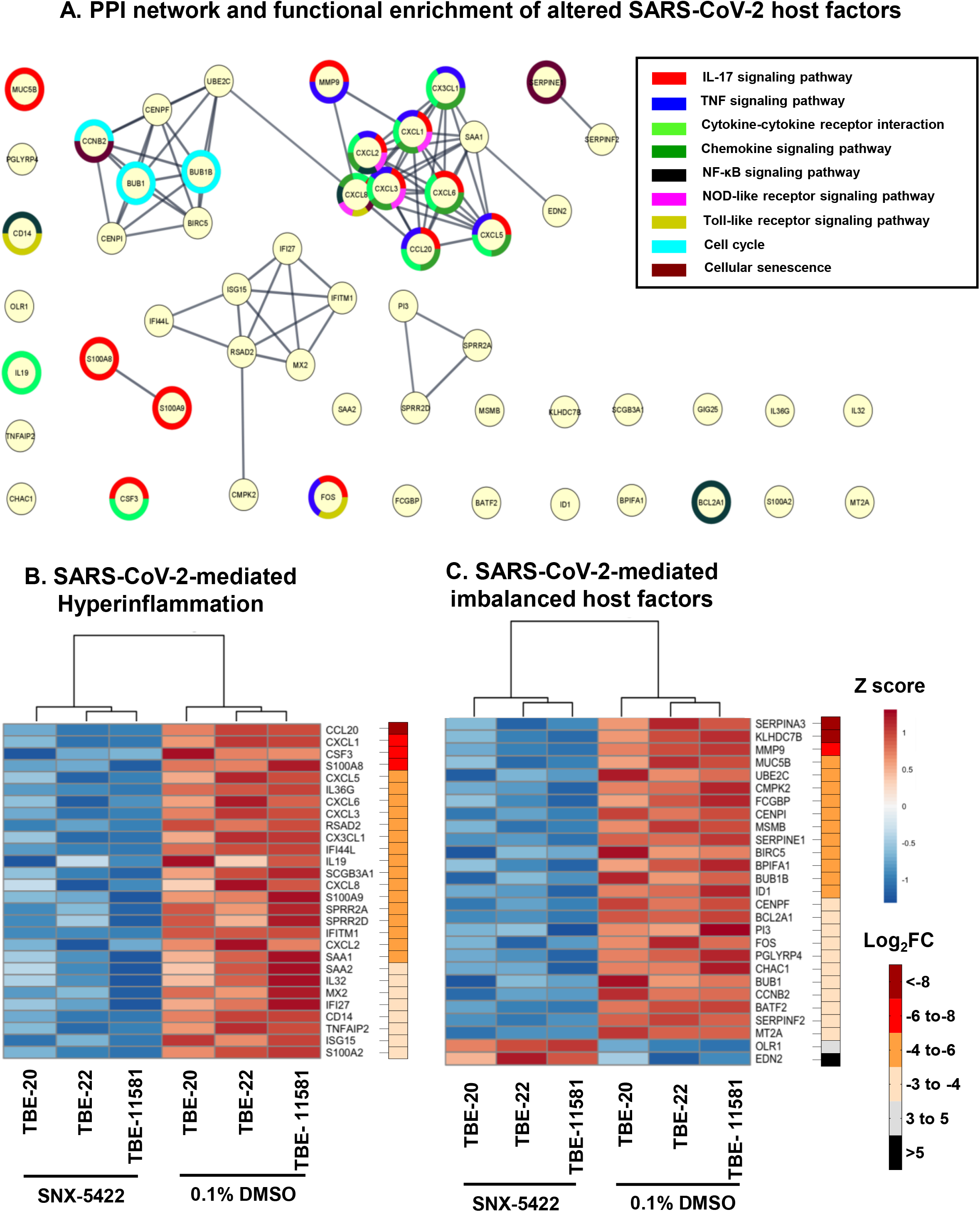
SNX-5422 treatment targets cellular pathways and genes involved in SARS-CoV-2 replication and disease progression. **(A)** Protein-protein interaction (PPI) network and functional enrichment of genes altered by SNX-5422 that are involved in SARS-CoV-2 replication and COVID-19 progression. Each node represents a protein and interactions with confidence score >0.9 are presented. Heat maps of DEGs following SNX-5422 treatment of human TBE cells, that have been identified to be **(B)** associated with SARS-CoV-2-mediated hyperinflammation and **(C)** imbalanced upon viral infection. Red represents relative upregulation of gene expression and blue represents relative downregulation of gene expression. Gene expression counts were transformed, and Z score normalized for heat map generation. Donors were clustered with complete linkage. Genes are arranged based on fold changes, as indicated by the fold change key.

### SNX-5422 regulates expression of host factors imbalanced upon SARS-CoV-2 infection

Recently, several host factors of SARS-CoV-2 replication have been identified (Gordon et al., 2020, Wang et al., 2021, Daniloski et al., 2021). Additionally, SARS-CoV-2 infection and COVID-19 progression have been associated with imbalanced expression of cellular pathways (Blanco-Melo et al., 2020) and heightened innate immune responses (Zhou et al., 2020a). Here, we aimed to investigate whether SNX-5422 treatment regulated expression of those previously identified SARS-CoV-2 host factors and cellular mechanisms imbalanced upon infection. Our data indicate that of the 55 genes altered by SNX-5422 treatment that were previously reported in the context of SARS-CoV-2 replication (Figure 3A), 27 genes had an imbalanced cellular expression, in virally infected cells (Figure 3C). Further categorization of these 55 genes revealed that the majority of these genes fall into 4 functional categories,-namely, regulators of innate immunity (*SERPINA3* (D’Alessandro et al., 2020)*, BPIFA1* (Liu et al., 2020), and *PGLYRP4* (Chandrashekar et al., 2020)), regulators of cell cycle (*UBE2C* (Bock and Ortea, 2020)*, CENPI* (Venkatakrishnan et al., 2020)*, BIRC5* (Xiong et al., 2020)*, CCNB2* (Xiong et al., 2020)*, BUB1B* (Ahmed et al., 2020)*, BUB1* (Ahmed et al., 2020) and *CENPF* (Gordon et al., 2020)), mucosal structural proteins (*MMP9* (Hazra et al., 2020) *and FCGBP* (Liu et al., 2020)) and regulators of blood coagulation (*SERPINA3* (D’Alessandro et al., 2020)*, SERPINE1 (McCord et al., 2020) and SERPINF2* (D’Alessandro et al., 2020)). Furthermore, cellular genes whose expression was impacted by SNX-5422 treatment, encoded for proteins that were previously shown to be associated with SARS-CoV-2 replication phases (*MMP9* (Hazra et al., 2020) and *SERPINE1 (McCord et al., 2020)*) or those interacting with SARS-CoV-2 structural and non-structural proteins (*BUB1B* (Ahmed et al., 2020)*, BUB1* (Ahmed et al., 2020), *CENPF* (Gordon et al., 2020) and *ID1* (Islam and Khan, 2020)). Taken together, our analysis indicate that SNX-5422 treatment interrupted cellular pathways and genes associated with replication of SARS-CoV-2.

Viruses rely on the biosynthetic machinery of the host for their replication. Therefore, an attractive strategy to attenuate viral propagation and mitigate poor disease outcomes is to target viral host-dependency factors. While the advantage of such an antiviral strategy is their efficacy against newly evolving viral variants and a high barrier towards development of drug resistant viral strains, such therapies might interfere with the normal cellular functioning. Hence, development of host-targeted therapeutics with very high selectivity index for suppressing viral replication yet having minimal off-target effect is crucial. Moreover, as post-exposure prophylactics are administered to individuals with no-to-mild symptoms, the safety bar for such a therapeutic modality will be higher than those administered in acutely infected patients. Hence, while host-directed drug candidates are currently fast-tracked into clinical trials as anti-SARS-CoV-2 therapeutics, characterizing the interactions of the drug candidates with host cell mechanisms will be absolutely critical to minimize adverse outcomes.

SARS-CoV-2 infection results in highly activated T cells (Chen and John Wherry, 2020), Th17 skewing (De Biasi et al., 2020), activation of the NF-κB signaling pathway (Huang et al., 2020b), and abnormal elevation of inflammatory cytokines (De Biasi et al., 2020), leading to hypercytokinemia and an imbalanced host immune response (Zhou et al., 2020a, Blanco-Melo et al., 2020). Hyperinflammation is a key determinant of the severity of COVID-19 outcomes (Mehta et al., 2020), where patients with higher levels of inflammatory cytokines have been associated with higher risk of lung injury and ARDS (Chen et al., 2020, Torres Acosta and Singer, 2020). Hence, targeted treatments, early during the course of infection, to prevent such cytokine storms have been proposed (Gallelli et al., 2020). We demonstrate that SNX-5422-treatment of human TBE cells downregulated key inflammatory pathways and genes associated with poor COVID-19 outcomes (Zhou et al., 2020a, Nienhold et al., 2020). Importantly, SNX-5422 treatment also altered expression of host responses reported to be imbalanced upon SARS-CoV-2 infection such as innate immune responses (Blanco-Melo et al., 2020), cell cycle regulation (Yuan et al., 2006), biosynthesis and metabolism of amino acids (Blasco et al., 2020). Moreover, SNX-5422 interrupted expression of cellular factors that directly or indirectly interact with the crucial steps of SARS-CoV-2 replication (McCord et al., 2020, Hazra et al., 2020) and with viral structural and non-structural proteins (Ahmed et al., 2020, Gordon et al., 2020, Islam and Khan, 2020).

In summary, our data support further testing and development of SNX-5422 as an oral post-exposure prophylactic or early-phase therapeutic for SARS-CoV-2 infection. Early administration of SNX-5422 will interfere with SARS-CoV-2 replication machinery and dampen virus-induced inflammation and severe disease outcomes, although human trials and experimental validation of such a mechanism in *ex vivo* human primary tissue sections or in animal models remains necessary. Additionally, the fact that SNX-5422 is orally bioavailable, and is clinically tested, makes this drug a quality candidate for an easy-to-administer and rapidly translatable post-exposure prophylactic therapy for SARS-CoV-2 infections. Finally, since SNX-5422 targets dependency factors that are utilized by multiple CoVs (Aldeyra Therapeutics Press Release, 2020, Emanuel et al., 2020, Iyad et al., 2020, Li et al., 2020, Li et al., 2004) this drug may have the potential to serve as a broad-spectrum ready-to-deploy therapeutic, which will be efficacious in not only ending the current pandemic, but also facilitating future pandemic preparedness against newly emerging, genetically diverse, CoV strains.

## ACKNOWLEDGEMENTS

This work was supported by a grant from Open Philanthropy (R.G and S.R.P) and COVID-19 research startup funds (S.R.P). The funders had no role in study design, data collection and interpretation, or the decision to submit the work for publication. The content is solely the responsibility of the authors. Work with live SARS-CoV-2 was performed under BSL-3 in the Duke Regional Biocontainment Laboratory (RBL), which received partial support for construction from the NIH/NIAD (UC6AI058607; G. D. Sempowski). Flow cytometry and fluorescent microscopy was performed at the DHVI Flow Cytometry Facility (Durham, NC) and Duke Light Microscopy Core facility (Durham, NC), respectively. Bulk-RNA sequencing was performed at the DHVI Viral Genetic Analysis core (Durham, NC). The following reagent was deposited by the Centers for Disease Control and Prevention and obtained through BEI Resources, NIAID, NIH: SARS-Related Coronavirus 2, Isolate USA-WA1/2020, NR-52281, and was provided by Dr. Gregory Sempowski. We thank Dr. Stacy Horner, Duke Molecular Genetics and Microbiology, for kindly providing Calu-3 (ATCC) cells.

## AUTHOR CONTRIBUTIONS

R.G., S.M.P., T.H., M.B. and S.R.P. designed the study and interpreted the data; R.G., J.J.T., F.K., S.N.L., J.S. and M.B. performed experiments and analyzed the data; V.R. performed the RNA-seq and pathway analysis; P.H. and T.H. provided SNX-5422 and consulted on the drug properties. S.M.P., T.H. and M.B. contributed important insights for the interpretation and discussion of the results. R.G. and S.R.P. drafted the manuscript. All authors read and approved the final manuscript.

## METHODS

### Cell cultures

Fully differentiated human TBE cells (EpiAirway™) from 3 independent donors with no reported respiratory disease or smoking history (Table S1) were obtained from MatTek (Ashland, MA). The cells were cultured at the air-liquid interface in 1 ml of AIR-100-MM culture medium (MatTek) in 6 well plates at 37°C in 5% CO_2_. Upon receipt of cells, the cultures were acclimated for 16-24hr prior to the start of experiments. Vero E6 cells (ATCC) were cultured in Dulbecco’s modified Eagle’s medium (DMEM) supplemented with 10% fetal bovine serum (FBS), 1X Penicillin/Streptomycin (Gibco), and 1X non-essential amino acid (NEAA) mixture (Gibco) and maintained at 37°C in 5% CO_2_. Calu-3 cells (ATCC) were cultured in Dulbecco’s modified Eagle’s medium (DMEM) supplemented with 20% fetal bovine serum (FBS), 25mM HEPES, 1X Penicillin/Streptomycin (Gibco), and maintained at 37°C in 5% CO_2_.

### Compounds

The small molecular inhibitor of Hsp90, SNX-5422, has been previously described (Fadden et al., 2010, Huang et al., 2009). The compound was synthesized in-house using previously described methods (Duan et al., 2012) and was characterized by proton nuclear magnetic resonance (NMR) and liquid chromatography/mass-spectrometry (LC/MS). The compound was solubilized in 100% DMSO to a 10mM stock concentration. Remdesivir (MedChemExpress LLC, USA) was reconstituted in 100% DMSO to a concentration of 10mM.

### SARS-CoV-2 propagation and titering

SARS-CoV-2 USA-WA1/2020 (BEI Resources; NR-52281) was propagated on Vero E6 cells at a multiplicity of infection (MOI) of 0.001 in virus diluent (DMEM supplemented with 2% FBS, 1X Penicillin/Streptomycin (Gibco), 1mM sodium pyruvate (Gibco) and 1X NEAA (Gibco)) at 37°C in 5% CO2. At day 4 post-infection (pi), cell supernatant containing the released virus was harvested, spun at 1500 rpm for 5 minutes, filtered through a 0.22μM sterile vacuum filtration system, aliquoted and stored at −80°C until further use.

Stock viral titer was determined by plaque assay. Briefly, 0.72 × 10^6^ Vero E6 cells were seeded in 6 well plates. The stock virus was serially diluted and incubated on cellular monolayer at 37°C in 5% CO2. After 1hr, virus was aspirated, and cells were overlayed with carboxy-methyl cellulose (CMC) containing media (0.6% CMC, MEM supplemented with 1X Penicillin/Streptomycin (Gibco), 2% FBS, 1mM sodium pyruvate (Gibco), 1X NEAA (Gibco), 0.3% sodium bicarbonate (Gibco), and 1X GlutaMAX (Gibco). After 4 days of incubation at 37°C in 5% CO2, plaque assays stained with 1% crystal violet in 10% neutral buffered formalin (NBF), and plaque forming unit/mL (Pfu/mL) was determined.

### SARS-CoV-2 infection and treatment

1.4 × 10^5^ Vero E6 or Calu-3 cells were seeded in 24 well plates. After 24 hrs, cells were incubated with the SARS-CoV-2 isolate at an MOI of 0.01-0.02 at 37°C and 5% CO_2_, with intermittent plate rocking. After 1hr, the virus was aspirated, cells were rinsed twice with 1X PBS and fresh maintenance medium containing dilutions of SNX-5422 or Remdesivir or 0.1% DMSO was added. The cells were incubated for 48hrs at 37°C and 5% CO_2_.

### Estimation of cellular cytotoxicity using flow cytometry

Suspensions of Vero E6 and Calu-3 cells were stained with LIVE/DEAD Fixable Aqua Dead Cell Stain kit (Thermo Fisher) according to manufacturer’s instructions. The stained cells were fixed using methanol-free 4% formaldehyde (Thermo Fisher) for 30 minutes and acquired on an LSRII flow cytometer (BD Biosciences) using BD FACS Diva software and analyzed with FlowJo software version 10.1 (Tree Star, Inc). LIVE/DEAD marker negative viable cells were selected from total cells after sequential selection of forward scatter and side scatter singlets.

### Infectious viral titer of supernatant

Cellular supernatant was collected from infected and drug-treated cells, 48hrs after infection. The supernatant was clarified by spinning at 1500rpm for 5 minutes and infectious viral titer was measured by plaque assay as described above.

### qRT-PCR for detecting cell-free viral RNA

SARS-CoV-2 RNA from cell supernatant was extracted using the QIAamp viral RNA mini kit (Qiagen). A two-step qRT-PCR was used to detect viral RNA released in the cell supernatant. In the first step, viral cDNA for the nucleocapsid (N) gene was generated using SuperScript III Reverse Transcriptase (Invitrogen) and N-reverse primer (5’-GAGGAACGAGAAGAGGCTTG-3’), following manufacturer’s instructions. In the second step, 7uL cDNA from step-1 was amplified using N gene forward primer (5’-CACATTGGCACCCGCAATC-3’), N gene reverse primer (5’-GAGGAACGAGAAGAGGCTTG-3’) and probe (5’-FAM-ACTTCCTCAAGGAACAACATTGCCA-QSY-3’) using Taqman mastermix (Thermo Fisher). The thermal cycling steps were: 50°C for 2 min, 95 °C for 10 min, and 40 cycles of 95 °C for 15 s and 60 °C for 1 min, and qPCR was performed on a Step-One-Plus real time PCR machine (Applied Biosystems) using the StepOne Software v2.3. Viral RNA copy number/mL supernatant was assessed using pCDNA3.1(+)-N-eGFP plasmid (GenScript) as standard.

### Immunofluorescence assay for detection of SARS-CoV-2 N protein

1.4 × 10^5^ Vero E6 cells were seeded in 4-well chamber slides (Corning). After 24 hrs, cells were incubated with SARS-CoV-2 USA-WA1/2020 strain at an MOI of 0.01 at 37°C and 5% CO_2_, with intermittent rocking. After 1hr, the virus was aspirated, cells were rinsed twice with 1X PBS and fresh maintenance medium containing dilutions of SNX-5422 or Remdesivir or 0.1% DMSO was added. After 48hrs, cells were fixed by submerging the chamber slides in 10% NBF for 2hrs. Fixed cells were washed three times with DPBS (Sigma) + 0.02% Triton X-100. For immunostaining, cells were permeabilized with DPBS+ 0.1% Triton-X-100 for 20 minutes and blocked in DPBS supplemented with 5% BSA, 0.02% Tween-20 and 10% donkey sera (Sigma) for 1hr. The cells were then incubated with anti-SARS-CoV-2 N protein antibody (Sinobiological; 1:100) for 2hrs at room temperature. After subsequent washing of the samples, the cells were treated with Alexa Fluor 488 donkey anti-rabbit secondary antibody (Invitrogen; 1:1000) for 1hr at room temperature. After further washing, the coverglasses were mounted on glass slides with ProLong Glass Antifade with NucBlue (Invitrogen) and sealed. Coverglasses were allowed to cure overnight prior to imaging using Axio Imager fluorescent microscope (Carl Zeiss). Images were captured with 20X objective, processed using Zeiss Zen Black software and proportion of SARS-CoV-2-infected cells were counted using Fiji, utilizing the analyze particle function.

### CC50 and IC50 calculations

CC50 and IC50 were calculated using Graph Pad Prism using curve fitting.

### Drug treatment and bulk RNA-sequencing of human TBE cells

Human TBE cells from 3 independent donors (Table S1) were treated with 1μM SNX-5422 or 0.1% DMSO added to the media on the basolateral side of the culture (Figure 2A), in three biological replicates. After 48hrs, cells were resuspended in TRIzol reagent (Thermo Fisher) and total RNA from the cells was extracted by phase separation with chloroform and subsequently using the RNeasy Mini Kit (Qiagen). RNA-Seq libraries were prepared using TruSeq RNA library Prep Kit v2 (Illumina, Inc. USA). Before pooling and sequencing, fragment length distribution and library quality were assessed on a TapeStation 2200 (Agilent Technologies), and the libraries were validated by Qubit Fluorometers (Thermo Fisher). All libraries were then pooled in a concentration at 4nM and sequenced on a NextSeq 500 Illumina sequencing platform system using NextSeq 500/550 High Output Kit v2.0 (150 cycles) (Illumina, Inc. USA).

### Analysis of the bulk RNA-seq data

RNA-Seq data was quality checked with FastQC (Andrews, 2010) and preprocessing was carried out using TrimGalore (Krueger) toolkit to trim low-quality bases and Illumina adapter sequences using default settings. Reads were aligned to the ENSEMBL Homo_sapiens.GRCh38.dna.primary_assembly genome using the ENSEMBL Homo_sapiens.GRCh38.100 transcript (Kersey et al., 2012) annotation file with STAR (Dobin et al., 2013) splice-aware RNA-seq alignment tool in paired mode allowing maximum multimapping of 3. Gene level counts were quantified using FeatureCounts (Liao et al., 2014) tool, counting unique features in non-stranded mode and retaining both gene ID, gene name, and gene biotype mapping from the ENSEMBL annotation file. Prior to differential expression analysis, count data was collapsed to donor level and genes for which mean raw count was at least 15 were kept. Normalization and differential expression were carried out with DESeq2 (Love et al., 2014) Bioconductor (Huber et al., 2015) package, utilizing the ‘apeglm’ Bioconductor package (Zhu et al., 2019) for log fold change shrinkage, in R statistical programming environment. The design formula was constructed to test for the effect of treatment while controlling for donor.

Principal component analysis (PCA) was performed with plotPCA in ‘DESeq2’ R package. PCA was done on variance stabilizing transformed (vst) count data and batch corrected with ‘limma’ package for genes with an average raw count of at least 15 across samples. Volcano plot with differentially expressed genes was generated using ‘EnhancedVolcano’ package in R, after filtering for described differential expression cutoffs. Heatmaps for differential expression of genes were generated using ‘pheatmap’ package in R. A vst was applied to count data, and batch corrected with ‘limma’ package, followed by Z-score normalization. Dot plots demonstrating upregulated and downregulated KEGG pathways were generated using ‘ClusterProfiler’ package in R using a “universe” of all human genes. Protein-protein interaction (PPI) network was constructed using STRING and Cytoscape (vs 3.8.2).

### Biocontainment and biosafety

Work with live SARS-CoV-2 isolate (USA-WA1/2020; BEI Resources NR-52281) was performed under Biosafety Level-3 (BSL-3) in the Duke Regional Biocontainment Laboratory.

### Material availability

This study did not generate new unique reagents. SNX-5422 can be obtained from T.H upon request.

### Data and code availability

The raw and processed RNA-seq data discussed in this publication have been deposited in NCBI’s Gene Expression Omnibus and are accessible through GEO Series accession number GSE166397 (https://www.ncbi.nlm.nih.gov/geo/query/acc.cgi?acc=GSE166397).

## SUPPLEMENTAL INFORMATION

**Figure S1. Sequencing read counts altered upon SNX-5422 treatment of human TBE cells**. Principal component analysis (PCA) of gene expression counts in human TBE cells treated with 0.1% DMSO (circles) or 1μM SNX-5422. (triangles). Each donor is indicated by a single color and 3 biological replicates from each donor are plotted. Red and blue oval symbols represent SNX-5422 and DMSO-treated clusters, respectively.

**Table S1.** Human Tracheobronchial cell (TBE) donor information.

**Table S2.** Differentially expressed genes (DEGs) used for cellular pathway overrepresentation analysis.

