## Supplemental figures and tables for "Oral Hsp90 inhibitor, SNX-5422, attenuates SARS-CoV-2 replication and dampens inflammation in airway cells"

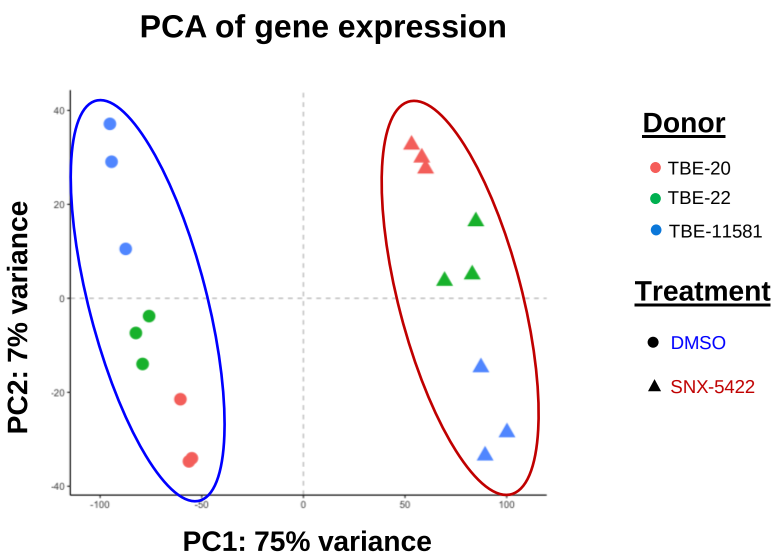


**Figure S1. Sequencing read counts altered upon SNX-5422 treatment of human TBE cells.** Principal component analysis (PCA) of gene expression counts in human TBE cells treated with 0.1% DMSO (circles) or 1µM SNX-5422. (triangles). Each donor is indicated by a single color and 3 biological replicates from each donor are plotted. Red and blue oval symbols represent SNX-5422 and DMSO-treated clusters, respectively.

**Table S1.** Human Tracheobronchial cell (TBE) donor information.

| Donor | Age | Sex[a] | Race[b] | Smoking status[c] | Lung disease[d] |
| --- | --- | --- | --- | --- | --- |
| TBE-20 | 13 | M | C | NS | N |
| TBE-22 | 35 | M | C | NS | N |
| TBE-11581 | 33 | F | C | NS | N |

[a]M: Male; F: Female
[b]C: Caucasian
[c]NS: Nonsmoker
[d]N: None identified

**Table S2.** Differentially expressed genes (DEGs) used for cellular pathway overrepresentation analysis.

| Gene Name | ENSEMBL ID | $Log_2FC$ (SNX-5422/DMSO) |
|---|---|---|
| SERPINA3 | ENSG00000196136 | -8.95 |
| SLC13A2 | ENSG00000007216 | -8.33 |
| CCL20 | ENSG00000115009 | -8.30 |
| FIBIN | ENSG00000176971 | -8.14 |
| KLHDC7B | ENSG00000130487 | -8.06 |
| MMP9 | ENSG00000100985 | -7.44 |
| HP | ENSG00000257017 | -7.08 |
| HLA-DOA | ENSG00000204252 | -6.98 |
| SLC26A4 | ENSG00000091137 | -6.85 |
| SLC5A1 | ENSG00000100170 | -6.72 |
| HLA-DMB | ENSG00000242574 | -6.36 |
| CXCL1 | ENSG00000163739 | -6.14 |
| CSF3 | ENSG00000108342 | -6.13 |
| CDC20 | ENSG00000117399 | -6.11 |
| S100A8 | ENSG00000143546 | -6.10 |
| SLC5A5 | ENSG00000105641 | -6.00 |
| TCIM | ENSG00000176907 | -5.97 |
| KLK12 | ENSG00000186474 | -5.94 |
| TNIP3 | ENSG00000050730 | -5.92 |
| DLGAP5 | ENSG00000126787 | -5.88 |
| TOP2A | ENSG00000131747 | -5.64 |
| CAPN13 | ENSG00000162949 | -5.52 |
| LRRC26 | ENSG00000184709 | -5.51 |
| CXCL5 | ENSG00000163735 | -5.49 |
| IL36G | ENSG00000136688 | -5.36 |
| MUC5B | ENSG00000117983 | -5.35 |
| CXCL6 | ENSG00000124875 | -5.32 |

| | | |
|---|---|---|
| UBE2C | ENSG00000175063 | -5.29 |
| CMPK2 | ENSG00000134326 | -5.27 |
| HLA-DRA | ENSG00000204287 | -5.24 |
| CXCL3 | ENSG00000163734 | -5.22 |
| RSAD2 | ENSG00000134321 | -5.20 |
| DUSP6 | ENSG00000139318 | -5.17 |
| ELN | ENSG00000049540 | -5.15 |
| CDC20B | ENSG00000164287 | -5.11 |
| CX3CL1 | ENSG00000006210 | -5.11 |
| IFI44L | ENSG00000137959 | -5.11 |
| IKBKE | ENSG00000263528 | -5.10 |
| HLA-DPB1 | ENSG00000223865 | -5.05 |
| CEP55 | ENSG00000138180 | -5.04 |
| FCGBP | ENSG00000275395 | -4.98 |
| C22orf15 | ENSG00000169314 | -4.97 |
| IL19 | ENSG00000142224 | -4.93 |
| CENPI | ENSG00000102384 | -4.92 |
| LTF | ENSG00000012223 | -4.91 |
| MSMB | ENSG00000263639 | -4.89 |
| DEPDC1 | ENSG00000024526 | -4.87 |
| RRM2 | ENSG00000171848 | -4.85 |
| SPRR1A | ENSG00000169474 | -4.83 |
| BPIFB1 | ENSG00000125999 | -4.81 |
| LCE3D | ENSG00000163202 | -4.80 |
| LDHD | ENSG00000166816 | -4.77 |
| CYP2F1 | ENSG00000197446 | -4.72 |
| CATIP | ENSG00000158428 | -4.70 |
| ANPEP | ENSG00000166825 | -4.69 |
| SCGB3A1 | ENSG00000161055 | -4.69 |

| | | |
|---|---|---|
| ALOX15B | ENSG00000179593 | -4.66 |
| CXCL8 | ENSG00000169429 | -4.65 |
| SERPINB4 | ENSG00000206073 | -4.64 |
| SKA3 | ENSG00000165480 | -4.62 |
| DOC2B | ENSG00000272636 | -4.60 |
| S100A9 | ENSG00000163220 | -4.59 |
| PHYHD1 | ENSG00000175287 | -4.58 |
| HLA-DQB1 | ENSG00000179344 | -4.52 |
| SERPINE1 | ENSG00000106366 | -4.43 |
| OR2I1P | ENSG00000237988 | -4.41 |
| MCF2L | ENSG00000126217 | -4.41 |
| BIRC5 | ENSG00000089685 | -4.37 |
| PYCR3 | ENSG00000104524 | -4.37 |
| NEK2 | ENSG00000117650 | -4.34 |
| KRT6B | ENSG00000185479 | -4.32 |
| MMP1 | ENSG00000196611 | -4.31 |
| BPIFA1 | ENSG00000198183 | -4.29 |
| S100A7 | ENSG00000143556 | -4.25 |
| TNC | ENSG00000041982 | -4.25 |
| BUB1B | ENSG00000156970 | -4.23 |
| ATP10A | ENSG00000206190 | -4.23 |
| SPRR2A | ENSG00000241794 | -4.21 |
| HLA-DRB1 | ENSG00000196126 | -4.20 |
| MYEOV | ENSG00000172927 | -4.20 |
| HLA-DQA1 | ENSG00000196735 | -4.19 |
| MAPK11 | ENSG00000185386 | -4.18 |
| SPRR2D | ENSG00000163216 | -4.17 |
| DAPL1 | ENSG00000163331 | -4.17 |
| MYCBPAP | ENSG00000136449 | -4.16 |

| | | |
|---|---|---|
| ID1 | ENSG00000125968 | -4.15 |
| PYCR1 | ENSG00000183010 | -4.14 |
| IFITM1 | ENSG00000185885 | -4.09 |
| CXCL2 | ENSG00000081041 | -4.07 |
| SAA1 | ENSG00000173432 | -4.07 |
| HLA-DPA1 | ENSG00000231389 | -4.07 |
| TRIB3 | ENSG00000101255 | -4.04 |
| IDO1 | ENSG00000131203 | -3.98 |
| MKI67 | ENSG00000148773 | -3.98 |
| DCDC2B | ENSG00000222046 | -3.98 |
| TTLL6 | ENSG00000170703 | -3.96 |
| LRRC74B | ENSG00000187905 | -3.96 |
| LENG9 | ENSG00000275183 | -3.95 |
| SAA2 | ENSG00000134339 | -3.95 |
| CYP2C18 | ENSG00000108242 | -3.92 |
| PDZK1IP1 | ENSG00000162366 | -3.90 |
| IL32 | ENSG00000008517 | -3.89 |
| KRT6C | ENSG00000170465 | -3.86 |
| KLK14 | ENSG00000129437 | -3.85 |
| ROPN1B | ENSG00000114547 | -3.85 |
| SDF2L1 | ENSG00000128228 | -3.85 |
| NCAPH | ENSG00000121152 | -3.84 |
| AMTN | ENSG00000187689 | -3.83 |
| LGALS7 | ENSG00000205076 | -3.81 |
| HLA-F | ENSG00000204642 | -3.78 |
| CENPF | ENSG00000117724 | -3.78 |
| SDSL | ENSG00000139410 | -3.77 |
| TGM2 | ENSG00000198959 | -3.75 |
| KLHDC9 | ENSG00000162755 | -3.73 |

| | | |
|---|---|---|
| LRRC71 | ENSG00000160838 | -3.72 |
| LRRC23 | ENSG00000010626 | -3.71 |
| KRT6A | ENSG00000205420 | -3.70 |
| NEURL3 | ENSG00000163121 | -3.69 |
| ABCC3 | ENSG00000108846 | -3.68 |
| BCL2A1 | ENSG00000140379 | -3.68 |
| DTL | ENSG00000143476 | -3.68 |
| CFB | ENSG00000243649 | -3.68 |
| UGT2B7 | ENSG00000171234 | -3.68 |
| HLA-DRB5 | ENSG00000198502 | -3.65 |
| SPAG8 | ENSG00000137098 | -3.63 |
| TTK | ENSG00000112742 | -3.63 |
| PCDHAC1 | ENSG00000248383 | -3.62 |
| BCL2L15 | ENSG00000188761 | -3.62 |
| CIITA | ENSG00000179583 | -3.61 |
| CNFN | ENSG00000105427 | -3.61 |
| SPRR2E | ENSG00000203785 | -3.60 |
| SAPCD2 | ENSG00000186193 | -3.60 |
| CABCOCO1 | ENSG00000183346 | -3.60 |
| KRT14 | ENSG00000186847 | -3.59 |
| SMIM6 | ENSG00000259120 | -3.58 |
| ASF1B | ENSG00000105011 | -3.58 |
| TMEM256 | ENSG00000205544 | -3.55 |
| GGACT | ENSG00000134864 | -3.54 |
| B9D1 | ENSG00000108641 | -3.54 |
| SLC4A4 | ENSG00000080493 | -3.54 |
| TMEM212 | ENSG00000186329 | -3.52 |
| TCF19 | ENSG00000137310 | -3.51 |
| DNAJC22 | ENSG00000178401 | -3.51 |

| | | |
|---|---|---|
| PI3 | ENSG00000124102 | -3.51 |
| FOS | ENSG00000170345 | -3.49 |
| SAA2-SAA4 | ENSG00000255071 | -3.47 |
| KCNJ15 | ENSG00000157551 | -3.47 |
| SLC34A2 | ENSG00000157765 | -3.45 |
| DERL3 | ENSG00000099958 | -3.45 |
| MFSD3 | ENSG00000167700 | -3.45 |
| MX2 | ENSG00000183486 | -3.45 |
| PGLYRP4 | ENSG00000163218 | -3.42 |
| FUOM | ENSG00000148803 | -3.42 |
| CHAC1 | ENSG00000128965 | -3.42 |
| IFI27 | ENSG00000165949 | -3.40 |
| BUB1 | ENSG00000169679 | -3.39 |
| ALDH1L2 | ENSG00000136010 | -3.39 |
| PRRX2 | ENSG00000167157 | -3.37 |
| RHEBL1 | ENSG00000167550 | -3.37 |
| CCNB2 | ENSG00000157456 | -3.37 |
| ERICH2 | ENSG00000204334 | -3.34 |
| RARRES1 | ENSG00000118849 | -3.33 |
| NUDT8 | ENSG00000167799 | -3.33 |
| KLK10 | ENSG00000129451 | -3.33 |
| MESP1 | ENSG00000166823 | -3.33 |
| PCK2 | ENSG00000100889 | -3.33 |
| TCTEX1D4 | ENSG00000188396 | -3.31 |
| GAS7 | ENSG00000007237 | -3.31 |
| LGALS1 | ENSG00000100097 | -3.29 |
| BPIFB2 | ENSG00000078898 | -3.29 |
| CD14 | ENSG00000170458 | -3.29 |
| PPOX | ENSG00000143224 | -3.29 |

| | | |
|---|---|---|
| G0S2 | ENSG00000123689 | -3.25 |
| MELK | ENSG00000165304 | -3.25 |
| FOXN4 | ENSG00000139445 | -3.24 |
| HLA-DMA | ENSG00000204257 | -3.20 |
| PSMB9 | ENSG00000240065 | -3.19 |
| TK1 | ENSG00000167900 | -3.19 |
| TNFAIP2 | ENSG00000185215 | -3.19 |
| ADAM8 | ENSG00000151651 | -3.17 |
| BATF2 | ENSG00000168062 | -3.16 |
| PEX11G | ENSG00000104883 | -3.16 |
| SNPH | ENSG00000101298 | -3.16 |
| CD200R1L | ENSG00000206531 | -3.15 |
| WDR97 | ENSG00000179698 | -3.13 |
| GLB1L | ENSG00000163521 | -3.13 |
| PPFIA3 | ENSG00000177380 | -3.13 |
| ETFBKMT | ENSG00000139160 | -3.12 |
| SLC6A20 | ENSG00000163817 | -3.12 |
| ISG15 | ENSG00000187608 | -3.11 |
| DLG4 | ENSG00000132535 | -3.10 |
| MEIG1 | ENSG00000197889 | -3.10 |
| TMEM92 | ENSG00000167105 | -3.10 |
| ERCC6L | ENSG00000186871 | -3.08 |
| INPP5J | ENSG00000185133 | -3.08 |
| IL27RA | ENSG00000104998 | -3.08 |
| SERPINF2 | ENSG00000167711 | -3.08 |
| MT1X | ENSG00000187193 | -3.07 |
| ETV4 | ENSG00000175832 | -3.07 |
| PPP1R42 | ENSG00000178125 | -3.07 |
| CFAP300 | ENSG00000137691 | -3.06 |

| | | |
|---|---|---|
| CCDC89 | ENSG00000179071 | -3.06 |
| MT2A | ENSG00000125148 | -3.06 |
| SVEP1 | ENSG00000165124 | -3.06 |
| CFAP73 | ENSG00000186710 | -3.06 |
| C10orf99 | ENSG00000188373 | -3.06 |
| POC1A | ENSG00000164087 | -3.05 |
| AZGP1 | ENSG00000160862 | -3.05 |
| KIFC1 | ENSG00000237649 | -3.05 |
| SLC52A3 | ENSG00000101276 | -3.04 |
| KNDC1 | ENSG00000171798 | -3.04 |
| XDH | ENSG00000158125 | -3.04 |
| EEF1AKMT4 | ENSG00000284753 | -3.03 |
| S100A2 | ENSG00000196754 | -3.03 |
| MMP10 | ENSG00000166670 | -3.03 |
| C16orf71 | ENSG00000166246 | -3.03 |
| ASRGL1 | ENSG00000162174 | -3.02 |
| EXOSC5 | ENSG00000077348 | -2.99 |
| DDIAS | ENSG00000165490 | -2.99 |
| ATP12A | ENSG00000075673 | -2.98 |
| IRF7 | ENSG00000185507 | -2.98 |
| C2orf73 | ENSG00000177994 | -2.98 |
| BCL2L14 | ENSG00000121380 | -2.97 |
| GSTA4 | ENSG00000170899 | -2.97 |
| TTLL10 | ENSG00000162571 | -2.96 |
| COL7A1 | ENSG00000114270 | -2.95 |
| SLCO4A1 | ENSG00000101187 | -2.95 |
| TPX2 | ENSG00000088325 | -2.94 |
| GNA14 | ENSG00000156049 | -2.93 |
| LRRC73 | ENSG00000204052 | -2.93 |

| | | |
|---|---|---|
| SLFN11 | ENSG00000172716 | -2.93 |
| DYNLRB2 | ENSG00000168589 | -2.92 |
| TESMIN | ENSG00000132749 | -2.92 |
| CROCC2 | ENSG00000226321 | -2.92 |
| SMIM35 | ENSG00000255274 | -2.92 |
| SLPI | ENSG00000124107 | -2.91 |
| TCN2 | ENSG00000185339 | -2.91 |
| TK2 | ENSG00000166548 | -2.91 |
| IGFBP6 | ENSG00000167779 | -2.89 |
| EGR1 | ENSG00000120738 | -2.89 |
| OASL | ENSG00000135114 | -2.88 |
| BPIFA2 | ENSG00000131050 | -2.88 |
| RPL22L1 | ENSG00000163584 | -2.88 |
| CCDC3 | ENSG00000151468 | -2.88 |
| ARHGAP40 | ENSG00000124143 | -2.88 |
| TMC5 | ENSG00000103534 | -2.87 |
| REEP2 | ENSG00000132563 | -2.86 |
| NUF2 | ENSG00000143228 | -2.86 |
| CDK1 | ENSG00000170312 | -2.86 |
| CAPN8 | ENSG00000203697 | -2.85 |
| RGS14 | ENSG00000169220 | -2.85 |
| KIF2C | ENSG00000142945 | -2.85 |
| TTC30A | ENSG00000197557 | -2.84 |
| CD164L2 | ENSG00000174950 | -2.84 |
| LGALS7B | ENSG00000178934 | -2.84 |
| C9orf24 | ENSG00000164972 | -2.84 |
| ELOVL6 | ENSG00000170522 | -2.84 |
| HMMR | ENSG00000072571 | -2.84 |
| CENPK | ENSG00000123219 | -2.83 |

| | | |
|---|---|---|
| PLEKHN1 | ENSG00000187583 | -2.83 |
| IQCH | ENSG00000103599 | -2.82 |
| MAD2L1 | ENSG00000164109 | -2.81 |
| SUSD4 | ENSG00000143502 | -2.80 |
| CCNA1 | ENSG00000133101 | -2.79 |
| UBE2T | ENSG00000077152 | -2.79 |
| CREB3L4 | ENSG00000143578 | -2.79 |
| CDCA5 | ENSG00000146670 | -2.76 |
| SCN4B | ENSG00000177098 | -2.75 |
| HLA-B | ENSG00000234745 | -2.75 |
| RSPH9 | ENSG00000172426 | -2.75 |
| SFXN2 | ENSG00000156398 | -2.74 |
| POP1 | ENSG00000104356 | -2.74 |
| FUT6 | ENSG00000156413 | -2.74 |
| CCT6B | ENSG00000132141 | -2.73 |
| SECTM1 | ENSG00000141574 | -2.73 |
| CENPW | ENSG00000203760 | -2.73 |
| SBSN | ENSG00000189001 | -2.72 |
| OPLAH | ENSG00000178814 | -2.72 |
| S100A12 | ENSG00000163221 | -2.72 |
| SRCIN1 | ENSG00000277363 | -2.71 |
| SERPINE2 | ENSG00000135919 | -2.71 |
| AK8 | ENSG00000165695 | -2.71 |
| CXCL14 | ENSG00000145824 | -2.71 |
| CRYM | ENSG00000103316 | -2.71 |
| RSPH14 | ENSG00000100218 | -2.70 |
| PLA2G4A | ENSG00000116711 | -2.70 |
| KIF20A | ENSG00000112984 | -2.70 |
| DNPH1 | ENSG00000112667 | -2.70 |

| | | |
|---|---|---|
| CKAP2L | ENSG00000169607 | -2.69 |
| TSTD1 | ENSG00000215845 | -2.69 |
| SERPINB3 | ENSG00000057149 | -2.69 |
| EMP1 | ENSG00000134531 | -2.68 |
| CALML5 | ENSG00000178372 | -2.68 |
| KLK13 | ENSG00000167759 | -2.68 |
| NPM3 | ENSG00000107833 | -2.67 |
| METTL1 | ENSG00000037897 | -2.67 |
| SPRR1B | ENSG00000169469 | -2.66 |
| CNPY4 | ENSG00000166997 | -2.66 |
| CKS1B | ENSG00000173207 | -2.66 |
| FBXO15 | ENSG00000141665 | -2.66 |
| PIP5KL1 | ENSG00000167103 | -2.66 |
| P3H4 | ENSG00000141696 | -2.66 |
| SMKR1 | ENSG00000240204 | -2.66 |
| TMEM107 | ENSG00000179029 | -2.65 |
| BCL3 | ENSG00000069399 | -2.65 |
| TMEM141 | ENSG00000244187 | -2.65 |
| PLA2G4F | ENSG00000168907 | -2.64 |
| KCTD14 | ENSG00000151364 | -2.64 |
| CNIH2 | ENSG00000174871 | -2.64 |
| C11orf1 | ENSG00000137720 | -2.63 |
| CCDC78 | ENSG00000162004 | -2.62 |
| PLGRKT | ENSG00000107020 | -2.62 |
| TLCD2 | ENSG00000185561 | -2.62 |
| GALK1 | ENSG00000108479 | -2.61 |
| LRRC46 | ENSG00000141294 | -2.61 |
| LRRC29 | ENSG00000125122 | -2.61 |
| RPL39L | ENSG00000163923 | -2.60 |

| | | |
|---|---|---|
| C1RL | ENSG00000139178 | -2.60 |
| ASS1 | ENSG00000130707 | -2.60 |
| PAQR4 | ENSG00000162073 | -2.59 |
| MMAB | ENSG00000139428 | -2.59 |
| KMO | ENSG00000117009 | -2.59 |
| NDUFAF3 | ENSG00000178057 | -2.58 |
| TFF3 | ENSG00000160180 | -2.58 |
| DEUP1 | ENSG00000165325 | -2.58 |
| RHOV | ENSG00000104140 | -2.58 |
| SAXO2 | ENSG00000188659 | -2.58 |
| MTHFD1L | ENSG00000120254 | -2.58 |
| SYT5 | ENSG00000129990 | -2.57 |
| GALNT14 | ENSG00000158089 | -2.57 |
| FOXM1 | ENSG00000111206 | -2.57 |
| PPP1R16B | ENSG00000101445 | -2.57 |
| RAC3 | ENSG00000169750 | -2.57 |
| IFIT1 | ENSG00000185745 | -2.57 |
| CRELD2 | ENSG00000184164 | -2.56 |
| FBF1 | ENSG00000188878 | -2.56 |
| CYP1B1 | ENSG00000138061 | -2.55 |
| CPXM2 | ENSG00000121898 | -2.54 |
| CRACR2B | ENSG00000177685 | -2.54 |
| ATP5MC1 | ENSG00000159199 | -2.53 |
| HEMK1 | ENSG00000114735 | -2.53 |
| BRCA2 | ENSG00000139618 | -2.53 |
| LIX1L | ENSG00000271601 | -2.53 |
| NUDT6 | ENSG00000170917 | -2.53 |
| CATSPERD | ENSG00000174898 | -2.52 |
| CENPM | ENSG00000100162 | -2.52 |

| | | |
|---|---|---|
| RNF183 | ENSG00000165188 | -2.52 |
| KATNAL2 | ENSG00000167216 | -2.52 |
| BIRC3 | ENSG00000023445 | -2.51 |
| MB | ENSG00000198125 | -2.51 |
| SLC2A6 | ENSG00000160326 | -2.51 |
| SPATA4 | ENSG00000150628 | -2.51 |
| CNGA4 | ENSG00000132259 | -2.50 |
| ADM2 | ENSG00000128165 | -2.50 |
| ADAM28 | ENSG00000042980 | -2.50 |
| CFAP74 | ENSG00000142609 | -2.50 |
| STEAP1 | ENSG00000164647 | -2.49 |
| FBXO4 | ENSG00000151876 | -2.49 |
| METTL27 | ENSG00000165171 | -2.49 |
| PREX1 | ENSG00000124126 | -2.48 |
| ACSL5 | ENSG00000197142 | -2.47 |
| CRACR2A | ENSG00000130038 | -2.47 |
| MUC1 | ENSG00000185499 | -2.47 |
| CD320 | ENSG00000167775 | -2.47 |
| HSD17B8 | ENSG00000204228 | -2.47 |
| MPV17L | ENSG00000156968 | -2.46 |
| FUT2 | ENSG00000176920 | -2.46 |
| BICDL2 | ENSG00000162069 | -2.46 |
| CHI3L1 | ENSG00000133048 | -2.45 |
| TGM5 | ENSG00000104055 | -2.45 |
| RWDD2B | ENSG00000156253 | -2.45 |
| MORN1 | ENSG00000116151 | -2.45 |
| ID2 | ENSG00000115738 | -2.45 |
| GMPPB | ENSG00000173540 | -2.44 |
| CDC42EP1 | ENSG00000128283 | -2.43 |

| | | |
|---|---|---|
| ATP8B2 | ENSG00000143515 | -2.43 |
| SMARCD3 | ENSG00000082014 | -2.42 |
| C1R | ENSG00000159403 | -2.42 |
| SPATA24 | ENSG00000170469 | -2.42 |
| RAD51AP1 | ENSG00000111247 | -2.41 |
| MORN5 | ENSG00000185681 | -2.41 |
| MPI | ENSG00000178802 | -2.41 |
| B3GNTL1 | ENSG00000175711 | -2.40 |
| IL36RN | ENSG00000136695 | -2.40 |
| DNAAF3 | ENSG00000167646 | -2.40 |
| AC007906.2 | ENSG00000277639 | -2.40 |
| TRIM14 | ENSG00000106785 | -2.40 |
| LYPD2 | ENSG00000197353 | -2.40 |
| MAPK15 | ENSG00000181085 | -2.39 |
| C1orf189 | ENSG00000163263 | -2.39 |
| SHMT2 | ENSG00000182199 | -2.38 |
| WDR93 | ENSG00000140527 | -2.38 |
| EMP3 | ENSG00000142227 | -2.38 |
| ASPG | ENSG00000166183 | -2.38 |
| PRADC1 | ENSG00000135617 | -2.37 |
| SPDEF | ENSG00000124664 | -2.37 |
| KRTDAP | ENSG00000188508 | -2.37 |
| SERPINB1 | ENSG00000021355 | -2.36 |
| C15orf48 | ENSG00000166920 | -2.36 |
| DGAT2 | ENSG00000062282 | -2.36 |
| AC055839.2 | ENSG00000286190 | -2.36 |
| ZG16B | ENSG00000162078 | -2.36 |
| EIF4EBP3 | ENSG00000243056 | -2.35 |
| IL34 | ENSG00000157368 | -2.35 |

| | | |
|---|---|---|
| ADGRE2 | ENSG00000127507 | -2.35 |
| SLC1A4 | ENSG00000115902 | -2.35 |
| STOML1 | ENSG00000067221 | -2.35 |
| RIBC2 | ENSG00000128408 | -2.35 |
| CYP4X1 | ENSG00000186377 | -2.35 |
| C1S | ENSG00000182326 | -2.34 |
| LY6D | ENSG00000167656 | -2.34 |
| C7orf57 | ENSG00000164746 | -2.34 |
| DHRS12 | ENSG00000102796 | -2.34 |
| VSTM2L | ENSG00000132821 | -2.34 |
| CD74 | ENSG00000019582 | -2.33 |
| TMEM121 | ENSG00000184986 | -2.32 |
| NFATC4 | ENSG00000100968 | -2.32 |
| PHGDH | ENSG00000092621 | -2.32 |
| ITPRIPL1 | ENSG00000198885 | -2.32 |
| ZCWPW1 | ENSG00000078487 | -2.31 |
| CXCL17 | ENSG00000189377 | -2.31 |
| OAS2 | ENSG00000111335 | -2.31 |
| FZD8 | ENSG00000177283 | -2.30 |
| SLC49A3 | ENSG00000169026 | -2.30 |
| PIGR | ENSG00000162896 | -2.30 |
| CFAP92 | ENSG00000114656 | -2.30 |
| RPL10A | ENSG00000198755 | -2.29 |
| HS3ST1 | ENSG00000002587 | -2.29 |
| DRC7 | ENSG00000159625 | -2.29 |
| FNDC11 | ENSG00000125531 | -2.29 |
| CDC25B | ENSG00000101224 | -2.29 |
| GJB6 | ENSG00000121742 | -2.28 |
| FAM20A | ENSG00000108950 | -2.28 |

| | | |
|---|---|---|
| C11orf97 | ENSG00000257057 | -2.28 |
| ALDH3B2 | ENSG00000132746 | -2.28 |
| LOXL1 | ENSG00000129038 | -2.28 |
| USP18 | ENSG00000184979 | -2.27 |
| SMPD2 | ENSG00000135587 | -2.27 |
| C10orf67 | ENSG00000179133 | -2.27 |
| CCDC159 | ENSG00000183401 | -2.26 |
| CCDC189 | ENSG00000196118 | -2.26 |
| BRCA1 | ENSG00000012048 | -2.26 |
| PCSK1N | ENSG00000102109 | -2.26 |
| SLC9A9 | ENSG00000181804 | -2.26 |
| MUC12 | ENSG00000205277 | -2.25 |
| LARGE2 | ENSG00000165905 | -2.25 |
| CFAP157 | ENSG00000160401 | -2.24 |
| ENO4 | ENSG00000188316 | -2.24 |
| TMEM45A | ENSG00000181458 | -2.23 |
| CLBA1 | ENSG00000140104 | -2.23 |
| PRR29 | ENSG00000224383 | -2.22 |
| CEACAM7 | ENSG00000007306 | -2.22 |
| PLB1 | ENSG00000163803 | -2.22 |
| TSNAXIP1 | ENSG00000102904 | -2.21 |
| SRD5A3 | ENSG00000128039 | -2.21 |
| WDR54 | ENSG00000005448 | -2.21 |
| NRBP2 | ENSG00000185189 | -2.21 |
| DOC2A | ENSG00000149927 | -2.21 |
| FAM229B | ENSG00000203778 | -2.20 |
| TTC29 | ENSG00000137473 | -2.20 |
| DHRS13 | ENSG00000167536 | -2.20 |
| ARPIN | ENSG00000242498 | -2.20 |

| | | |
|---|---|---|
| C15orf65 | ENSG00000261652 | -2.20 |
| CCDC96 | ENSG00000173013 | -2.20 |
| HAUS5 | ENSG00000249115 | -2.19 |
| IFI6 | ENSG00000126709 | -2.19 |
| TCTE1 | ENSG00000146221 | -2.19 |
| LRRIQ3 | ENSG00000162620 | -2.19 |
| KIAA0040 | ENSG00000235750 | -2.19 |
| XAF1 | ENSG00000132530 | -2.19 |
| NAT14 | ENSG00000090971 | -2.19 |
| SCGB1A1 | ENSG00000149021 | -2.18 |
| CFAP221 | ENSG00000163075 | -2.18 |
| KIF18A | ENSG00000121621 | -2.18 |
| ARMH1 | ENSG00000198520 | -2.18 |
| COL9A2 | ENSG00000049089 | -2.17 |
| GCDH | ENSG00000105607 | -2.17 |
| CFAP57 | ENSG00000243710 | -2.17 |
| CCDC121 | ENSG00000176714 | -2.17 |
| UROS | ENSG00000188690 | -2.17 |
| RPL12 | ENSG00000197958 | -2.17 |
| FKBP10 | ENSG00000141756 | -2.16 |
| TMEM178B | ENSG00000261115 | -2.16 |
| CCDC61 | ENSG00000104983 | -2.16 |
| FAM166B | ENSG00000215187 | -2.15 |
| PAXX | ENSG00000148362 | -2.15 |
| ADPRHL1 | ENSG00000153531 | -2.15 |
| GPR68 | ENSG00000119714 | -2.15 |
| TRPT1 | ENSG00000149743 | -2.15 |
| TTC30B | ENSG00000196659 | -2.14 |
| NLRC5 | ENSG00000140853 | -2.14 |

| | | |
|---|---|---|
| FAM183A | ENSG00000186973 | -2.13 |
| RIBC1 | ENSG00000158423 | -2.13 |
| TIMP4 | ENSG00000157150 | -2.13 |
| KYNU | ENSG00000115919 | -2.13 |
| SPAG17 | ENSG00000155761 | -2.13 |
| CYBA | ENSG00000051523 | -2.13 |
| IRAK3 | ENSG00000090376 | -2.13 |
| H6PD | ENSG00000049239 | -2.13 |
| UBXN11 | ENSG00000158062 | -2.12 |
| LIF | ENSG00000128342 | -2.12 |
| ASTN2 | ENSG00000148219 | -2.12 |
| AKAP3 | ENSG00000111254 | -2.12 |
| PRRT3 | ENSG00000163704 | -2.12 |
| COPZ2 | ENSG00000005243 | -2.11 |
| LMNB1 | ENSG00000113368 | -2.11 |
| OAS3 | ENSG00000111331 | -2.11 |
| NOXA1 | ENSG00000188747 | -2.11 |
| ACYP1 | ENSG00000119640 | -2.11 |
| TMEM190 | ENSG00000160472 | -2.11 |
| MORN3 | ENSG00000139714 | -2.11 |
| STC2 | ENSG00000113739 | -2.11 |
| SLC19A1 | ENSG00000173638 | -2.10 |
| CFAP161 | ENSG00000156206 | -2.10 |
| IFI44 | ENSG00000137965 | -2.10 |
| POLR3GL | ENSG00000121851 | -2.10 |
| KDELR3 | ENSG00000100196 | -2.10 |
| IFTAP | ENSG00000166352 | -2.09 |
| IQCD | ENSG00000166578 | -2.09 |
| MAFF | ENSG00000185022 | -2.09 |

| | | |
|---|---|---|
| PIAS3 | ENSG00000131788 | -2.09 |
| LIPT2 | ENSG00000175536 | -2.09 |
| BBOX1 | ENSG00000129151 | -2.09 |
| CCDC74B | ENSG00000152076 | -2.09 |
| LRMDA | ENSG00000148655 | -2.09 |
| WFDC2 | ENSG00000101443 | -2.08 |
| WDR38 | ENSG00000136918 | -2.08 |
| TEDC1 | ENSG00000185347 | -2.08 |
| BPHL | ENSG00000137274 | -2.08 |
| MICOS13 | ENSG00000174917 | -2.07 |
| UHRF1 | ENSG00000276043 | -2.07 |
| EPHX2 | ENSG00000120915 | -2.07 |
| KLK7 | ENSG00000169035 | -2.06 |
| LIN7B | ENSG00000104863 | -2.06 |
| IRAK2 | ENSG00000134070 | -2.06 |
| CCL22 | ENSG00000102962 | -2.06 |
| NAV1 | ENSG00000134369 | -2.05 |
| TNFAIP3 | ENSG00000118503 | -2.05 |
| PLCB2 | ENSG00000137841 | -2.05 |
| ZC3H12A | ENSG00000163874 | -2.05 |
| PHLDA2 | ENSG00000181649 | -2.04 |
| FAM216A | ENSG00000204856 | -2.03 |
| STMN3 | ENSG00000197457 | -2.03 |
| MYCL | ENSG00000116990 | -2.03 |
| SUSD2 | ENSG00000099994 | -2.03 |
| HMBS | ENSG00000256269 | -2.03 |
| CKMT1B | ENSG00000237289 | -2.02 |
| EEF2KMT | ENSG00000118894 | -2.02 |
| RELB | ENSG00000104856 | -2.02 |

| | | |
|---|---|---|
| PDE9A | ENSG00000160191 | -2.02 |
| PGBD5 | ENSG00000177614 | -2.02 |
| KCNJ5 | ENSG00000120457 | -2.02 |
| PTGS2 | ENSG00000073756 | -2.02 |
| ZNF648 | ENSG00000179930 | -2.02 |
| PNMA8A | ENSG00000182013 | -2.01 |
| DNAJC12 | ENSG00000108176 | -2.01 |
| SEC11C | ENSG00000166562 | -2.01 |
| TMEM220 | ENSG00000187824 | -2.01 |
| KIF6 | ENSG00000164627 | -2.01 |
| SDHAF4 | ENSG00000154079 | -2.00 |
| ETV7 | ENSG00000010030 | -2.00 |
| TLR2 | ENSG00000137462 | -2.00 |
| RNF217 | ENSG00000146373 | 2.00 |
| CPM | ENSG00000135678 | 2.00 |
| XRN1 | ENSG00000114127 | 2.01 |
| CLIP3 | ENSG00000105270 | 2.01 |
| SELENOP | ENSG00000250722 | 2.02 |
| CALD1 | ENSG00000122786 | 2.02 |
| HCLS1 | ENSG00000180353 | 2.03 |
| CNTNAP3B | ENSG00000154529 | 2.03 |
| LATS2 | ENSG00000150457 | 2.04 |
| PANX2 | ENSG00000073150 | 2.04 |
| HELZ | ENSG00000198265 | 2.04 |
| PHTF2 | ENSG00000006576 | 2.05 |
| CCNG1 | ENSG00000113328 | 2.05 |
| PDE5A | ENSG00000138735 | 2.05 |
| ITSN1 | ENSG00000205726 | 2.05 |
| MAF | ENSG00000178573 | 2.06 |

| | | |
|---|---|---|
| DSEL | ENSG00000171451 | 2.07 |
| KCNJ14 | ENSG00000182324 | 2.07 |
| SIK1B | ENSG00000275993 | 2.07 |
| ZSCAN31 | ENSG00000235109 | 2.07 |
| CAV1 | ENSG00000105974 | 2.09 |
| LAMA4 | ENSG00000112769 | 2.10 |
| FAM13A | ENSG00000138640 | 2.10 |
| KDM6A | ENSG00000147050 | 2.10 |
| RRM2B | ENSG00000048392 | 2.10 |
| FOSL2 | ENSG00000075426 | 2.10 |
| C1QTNF6 | ENSG00000133466 | 2.11 |
| NRG1 | ENSG00000157168 | 2.11 |
| SEMA3E | ENSG00000170381 | 2.11 |
| UCP3 | ENSG00000175564 | 2.11 |
| CHML | ENSG00000203668 | 2.12 |
| DNAJB4 | ENSG00000162616 | 2.12 |
| ATP13A3 | ENSG00000133657 | 2.13 |
| ARHGAP22 | ENSG00000128805 | 2.13 |
| KIAA0513 | ENSG00000135709 | 2.13 |
| RAB30 | ENSG00000137502 | 2.13 |
| OTUD7B | ENSG00000264522 | 2.14 |
| DENND5B | ENSG00000170456 | 2.14 |
| SLC39A1 | ENSG00000143570 | 2.14 |
| SLAMF7 | ENSG00000026751 | 2.14 |
| TUFT1 | ENSG00000143367 | 2.15 |
| RETREG1 | ENSG00000154153 | 2.15 |
| PTPN14 | ENSG00000152104 | 2.15 |
| CLDN15 | ENSG00000106404 | 2.15 |
| RASD1 | ENSG00000108551 | 2.15 |

| | | |
|---|---|---|
| TMEM131 | ENSG00000075568 | 2.16 |
| RGS2 | ENSG00000116741 | 2.17 |
| TMEM64 | ENSG00000180694 | 2.17 |
| EXPH5 | ENSG00000110723 | 2.17 |
| SCHIP1 | ENSG00000151967 | 2.17 |
| SYNE2 | ENSG00000054654 | 2.18 |
| RALGPS2 | ENSG00000116191 | 2.18 |
| RUSC2 | ENSG00000198853 | 2.18 |
| NPBWR1 | ENSG00000288611 | 2.18 |
| AAK1 | ENSG00000115977 | 2.19 |
| ANKRD2 | ENSG00000165887 | 2.19 |
| PDK4 | ENSG00000004799 | 2.19 |
| CDK14 | ENSG00000058091 | 2.19 |
| DPEP2 | ENSG00000167261 | 2.20 |
| AGL | ENSG00000162688 | 2.20 |
| TTC28 | ENSG00000100154 | 2.21 |
| PIK3CA | ENSG00000121879 | 2.21 |
| CASTOR2 | ENSG00000274070 | 2.21 |
| PALLD | ENSG00000129116 | 2.21 |
| RFLNB | ENSG00000183688 | 2.21 |
| TGFBR3 | ENSG00000069702 | 2.22 |
| PER2 | ENSG00000132326 | 2.22 |
| CDK6 | ENSG00000105810 | 2.23 |
| ITGB2 | ENSG00000160255 | 2.23 |
| DSG2 | ENSG00000046604 | 2.24 |
| AQP4 | ENSG00000171885 | 2.25 |
| WNT3A | ENSG00000154342 | 2.26 |
| TIMP2 | ENSG00000035862 | 2.26 |
| USP31 | ENSG00000103404 | 2.26 |

| | | |
|---|---|---|
| CNTN3 | ENSG00000113805 | 2.26 |
| HSPA4L | ENSG00000164070 | 2.26 |
| SLC12A6 | ENSG00000140199 | 2.27 |
| TMEM156 | ENSG00000121895 | 2.27 |
| TGFB2 | ENSG00000092969 | 2.27 |
| ARHGAP29 | ENSG00000137962 | 2.27 |
| TGFB3 | ENSG00000119699 | 2.28 |
| DNAH17 | ENSG00000187775 | 2.28 |
| SESN3 | ENSG00000149212 | 2.28 |
| PGAP1 | ENSG00000197121 | 2.28 |
| SCARA3 | ENSG00000168077 | 2.28 |
| HOMER1 | ENSG00000152413 | 2.29 |
| HSPA1B | ENSG00000204388 | 2.29 |
| GRAMD1B | ENSG00000023171 | 2.29 |
| GREB1L | ENSG00000141449 | 2.30 |
| FMO2 | ENSG00000094963 | 2.30 |
| AKT3 | ENSG00000117020 | 2.30 |
| AL031777.2 | ENSG00000282988 | 2.30 |
| TNS1 | ENSG00000079308 | 2.31 |
| HEG1 | ENSG00000173706 | 2.31 |
| LAMA1 | ENSG00000101680 | 2.31 |
| ANKRD18B | ENSG00000230453 | 2.31 |
| FILIP1 | ENSG00000118407 | 2.31 |
| AFF1 | ENSG00000172493 | 2.32 |
| AL157935.2 | ENSG00000257524 | 2.32 |
| SRGAP1 | ENSG00000196935 | 2.33 |
| PROS1 | ENSG00000184500 | 2.33 |
| TEAD1 | ENSG00000187079 | 2.33 |
| ERP27 | ENSG00000139055 | 2.34 |

| | | |
|---|---|---|
| CAVIN2 | ENSG00000168497 | 2.34 |
| ADAM12 | ENSG00000148848 | 2.34 |
| ITGB3 | ENSG00000259207 | 2.34 |
| ACER2 | ENSG00000177076 | 2.34 |
| ABCA1 | ENSG00000165029 | 2.36 |
| RBMS3 | ENSG00000144642 | 2.36 |
| PDE7B | ENSG00000171408 | 2.36 |
| TMEM169 | ENSG00000163449 | 2.36 |
| FRRS1 | ENSG00000156869 | 2.37 |
| CLSPN | ENSG00000092853 | 2.37 |
| PDZD2 | ENSG00000133401 | 2.38 |
| SEMA3B | ENSG00000012171 | 2.38 |
| BCO1 | ENSG00000135697 | 2.38 |
| GCOM1 | ENSG00000137878 | 2.39 |
| CEMIP2 | ENSG00000135048 | 2.39 |
| SPOCD1 | ENSG00000134668 | 2.39 |
| FAM189A2 | ENSG00000135063 | 2.40 |
| BLNK | ENSG00000095585 | 2.40 |
| UPK3B | ENSG00000243566 | 2.41 |
| PLAC9 | ENSG00000189129 | 2.42 |
| ARHGEF26 | ENSG00000114790 | 2.43 |
| SCUBE3 | ENSG00000146197 | 2.44 |
| SLC16A7 | ENSG00000118596 | 2.44 |
| CAP2 | ENSG00000112186 | 2.45 |
| RBM20 | ENSG00000203867 | 2.45 |
| FAT1 | ENSG00000083857 | 2.45 |
| ABCG1 | ENSG00000160179 | 2.45 |
| DEPP1 | ENSG00000165507 | 2.46 |
| S1PR1 | ENSG00000170989 | 2.46 |

| | | |
|---|---|---|
| PWWP3B | ENSG00000157502 | 2.47 |
| PTCHD4 | ENSG00000244694 | 2.48 |
| SLC1A1 | ENSG00000106688 | 2.48 |
| FAXC | ENSG00000146267 | 2.48 |
| SULT1C2 | ENSG00000198203 | 2.48 |
| PTGER4 | ENSG00000171522 | 2.49 |
| FLG | ENSG00000143631 | 2.49 |
| DSC3 | ENSG00000134762 | 2.50 |
| PTPRQ | ENSG00000139304 | 2.50 |
| FRY | ENSG00000073910 | 2.50 |
| PLEKHA1 | ENSG00000107679 | 2.51 |
| GALR2 | ENSG00000182687 | 2.52 |
| ABCA6 | ENSG00000154262 | 2.52 |
| SEMA5A | ENSG00000112902 | 2.52 |
| CYBRD1 | ENSG00000071967 | 2.53 |
| OXTR | ENSG00000180914 | 2.53 |
| SYNPO2 | ENSG00000172403 | 2.55 |
| AATK | ENSG00000181409 | 2.55 |
| ZBED2 | ENSG00000177494 | 2.55 |
| CYP2U1 | ENSG00000155016 | 2.57 |
| DAB2 | ENSG00000153071 | 2.57 |
| MSRB3 | ENSG00000174099 | 2.58 |
| GBP1 | ENSG00000117228 | 2.59 |
| RASSF6 | ENSG00000169435 | 2.59 |
| AXL | ENSG00000167601 | 2.59 |
| INKA2 | ENSG00000197852 | 2.60 |
| CRIM1 | ENSG00000150938 | 2.60 |
| VSIG10 | ENSG00000176834 | 2.60 |
| ATP6V1B1 | ENSG00000116039 | 2.61 |

| | | |
|---|---|---|
| FERMT2 | ENSG00000073712 | 2.63 |
| GLCCI1 | ENSG00000106415 | 2.63 |
| HSPA1A | ENSG00000204389 | 2.65 |
| SMPDL3A | ENSG00000172594 | 2.65 |
| LHFPL6 | ENSG00000183722 | 2.65 |
| PRKAB2 | ENSG00000131791 | 2.65 |
| CRYAB | ENSG00000109846 | 2.66 |
| RARB | ENSG00000077092 | 2.67 |
| SORL1 | ENSG00000137642 | 2.67 |
| SYDE2 | ENSG00000097096 | 2.68 |
| MAP3K14 | ENSG00000006062 | 2.70 |
| HMGB3 | ENSG00000029993 | 2.71 |
| NNAT | ENSG00000053438 | 2.71 |
| PAMR1 | ENSG00000149090 | 2.72 |
| SPARC | ENSG00000113140 | 2.72 |
| DOCK8 | ENSG00000107099 | 2.73 |
| CMYA5 | ENSG00000164309 | 2.73 |
| CORO6 | ENSG00000167549 | 2.74 |
| WNT3 | ENSG00000108379 | 2.74 |
| N4BP2L1 | ENSG00000139597 | 2.75 |
| ELOVL7 | ENSG00000164181 | 2.75 |
| SRPX2 | ENSG00000102359 | 2.77 |
| S1PR4 | ENSG00000125910 | 2.77 |
| HSD17B11 | ENSG00000198189 | 2.77 |
| DOCK3 | ENSG00000088538 | 2.78 |
| RAB3B | ENSG00000169213 | 2.79 |
| FBLN2 | ENSG00000163520 | 2.80 |
| SSUH2 | ENSG00000125046 | 2.80 |
| ARG2 | ENSG00000081181 | 2.83 |

| | | |
|---|---|---|
| CLCA4 | ENSG00000016602 | 2.84 |
| STARD13 | ENSG00000133121 | 2.84 |
| FABP3 | ENSG00000121769 | 2.85 |
| ANKRD33B | ENSG00000164236 | 2.85 |
| CYP4F3 | ENSG00000186529 | 2.86 |
| SCIN | ENSG00000006747 | 2.87 |
| NCR3LG1 | ENSG00000188211 | 2.88 |
| OLFML2A | ENSG00000185585 | 2.88 |
| NPNT | ENSG00000168743 | 2.90 |
| PPP1R12B | ENSG00000077157 | 2.95 |
| HSD17B6 | ENSG00000025423 | 2.96 |
| MAP2 | ENSG00000078018 | 2.99 |
| LRAT | ENSG00000121207 | 3.02 |
| METTL7A | ENSG00000185432 | 3.02 |
| SLC40A1 | ENSG00000138449 | 3.07 |
| PIEZO2 | ENSG00000154864 | 3.07 |
| CLDN16 | ENSG00000113946 | 3.08 |
| SULT1E1 | ENSG00000109193 | 3.08 |
| INPP5D | ENSG00000168918 | 3.12 |
| ALOX12 | ENSG00000108839 | 3.14 |
| POU2F3 | ENSG00000137709 | 3.16 |
| PCDH19 | ENSG00000165194 | 3.18 |
| VGLL3 | ENSG00000206538 | 3.19 |
| COL4A4 | ENSG00000081052 | 3.20 |
| MPP2 | ENSG00000108852 | 3.20 |
| CLEC2B | ENSG00000110852 | 3.21 |
| ARL14 | ENSG00000179674 | 3.22 |
| SH2D1B | ENSG00000198574 | 3.23 |
| PCDH9 | ENSG00000184226 | 3.24 |

| | | |
|---|---|---|
| GLIPR1 | ENSG00000139278 | 3.25 |
| CEL | ENSG00000170835 | 3.27 |
| TNNC1 | ENSG00000114854 | 3.29 |
| IL12A | ENSG00000168811 | 3.30 |
| B4GALNT2 | ENSG00000167080 | 3.30 |
| MYO16 | ENSG00000041515 | 3.32 |
| AOX1 | ENSG00000138356 | 3.37 |
| ANK1 | ENSG00000029534 | 3.44 |
| EDIL3 | ENSG00000164176 | 3.45 |
| IL1RL1 | ENSG00000115602 | 3.46 |
| HOXA5 | ENSG00000106004 | 3.50 |
| HSD17B13 | ENSG00000170509 | 3.53 |
| P3H2 | ENSG00000090530 | 3.53 |
| F5 | ENSG00000198734 | 3.58 |
| CILP | ENSG00000138615 | 3.61 |
| PGF | ENSG00000119630 | 3.61 |
| IL31RA | ENSG00000164509 | 3.62 |
| SLC10A5 | ENSG00000253598 | 3.65 |
| GPRASP1 | ENSG00000198932 | 3.68 |
| UGT2A1 | ENSG00000173610 | 3.68 |
| BCL2 | ENSG00000171791 | 3.69 |
| OVCH2 | ENSG00000183378 | 3.72 |
| WFIKKN1 | ENSG00000127578 | 3.76 |
| HOXA1 | ENSG00000105991 | 3.76 |
| EPHX4 | ENSG00000172031 | 3.83 |
| PLEKHO1 | ENSG00000023902 | 3.85 |
| PDGFD | ENSG00000170962 | 3.85 |
| LOXL4 | ENSG00000138131 | 3.86 |
| GPR176 | ENSG00000166073 | 3.89 |

| | | |
|---|---|---|
| LUM | ENSG00000139329 | 3.93 |
| FGD3 | ENSG00000127084 | 4.00 |
| SLC30A10 | ENSG00000196660 | 4.05 |
| COL8A1 | ENSG00000144810 | 4.15 |
| SEMA6D | ENSG00000137872 | 4.16 |
| FAT4 | ENSG00000196159 | 4.24 |
| DPP4 | ENSG00000197635 | 4.27 |
| NHSL2 | ENSG00000204131 | 4.28 |
| CYP2W1 | ENSG00000073067 | 4.42 |
| STON1 | ENSG00000243244 | 4.47 |
| OLR1 | ENSG00000173391 | 4.53 |
| PEG10 | ENSG00000242265 | 4.54 |
| MYL9 | ENSG00000101335 | 4.63 |
| LYPD1 | ENSG00000150551 | 4.64 |
| PRH2 | ENSG00000134551 | 4.73 |
| PSG5 | ENSG00000204941 | 4.78 |
| CYP26B1 | ENSG00000003137 | 4.84 |
| NEXN | ENSG00000162614 | 5.01 |
| EDN2 | ENSG00000127129 | 5.16 |
| PNPLA1 | ENSG00000180316 | 5.26 |
| CDH5 | ENSG00000179776 | 6.03 |
| PTPRR | ENSG00000153233 | 6.08 |
| SPOCK2 | ENSG00000107742 | 6.09 |
| TMEM255A | ENSG00000125355 | 6.14 |
